# Scalable genotyping in fixed transcriptomes resolves clonal heterogeneity via single-cell sequencing

**DOI:** 10.64898/2026.04.11.717967

**Authors:** Sydney B. Blattman, Nabih Maslah, Austin A. Varela, Karolis Kumpaitis, Benan Nalbant, Catherine Snopkowski, Marisa Mariani, Laura C. Kida, Meril Takizawa, Nalin Ratnayeke, Kenny Kwok Hei Yu, Sunjay Fernandes, Nima Mousavi, Erik Borgstrom, Derek Vallejo, Lorita Boghospor, Ruijiao Xin, Marco Mignardi, Snow Wu, Nicholas Scarlott, Loruhama Delgado-Rivera, Poornasree Kumar, Sreenath Krishnan, Stéphane Giraudier, Jean-Jacques Kiladjian, Brooke E. Howitt, Andrew Kohlway, Paul Lund, Dana Pe’er, Ronan Chaligné, Caleb A. Lareau

**Author notes:** These authors contributed equally (Sydney B. Blattman, Nabih Maslah, Austin A. Varela).

## Abstract

Despite the promise of single-cell transcriptomics for understanding cell states in heterogeneous populations, widely used platforms have limited ability to link transcriptional states to somatic mutations within the same cells. Here, we introduce Genotyping in Fixed Transcriptomes (GIFT) for the simultaneous detection of large numbers of targeted genetic variants with whole transcriptome profiles in single cells. The core innovation of GIFT is a rationally designed gapfilling reaction between adjacent single-stranded DNA (ssDNA) probes that barcodes native transcript sequence to enable highly-specific targeted mutation detection. GIFT achieves greater than 99% genotyping accuracy and flexible capture of hundreds of mutations per cell, including in formalin-fixed, paraffin-embedded (FFPE) tissue, enabling clonal lineage tracing in heterogeneous settings. We demonstrate the unique scalability of GIFT by profiling more than 700,000 cells from 35 donors with myeloproliferative neoplasms (MPN), revealing mutation-dependent hematopoietic responses to systemic inflammation associated with the characteristic *JAK2*V617 mutation, including an allelic dose gradient of interferon-associated transcriptional programs and priming of hematopoietic stem cells that develop into divergent disease states. The technical advantages of GIFT enable direct resolution of genotype-to-phenotype relationships via clonal tracing with comprehensive cell-state measurements at single-cell resolution.

## Introduction

During normal development and aging, somatic DNA mutations accumulate at each cell division. These mutations can influence cellular differentiation, expansion, adaptation, and survival^1^. Decades of next-generation sequencing have revealed that most tumors harbor multiple oncogenic driver mutations as well as dozens to hundreds of passenger mutations^2^. Comprehensive mutation profiles can be used to infer somatic evolution by revealing relationships between subclones, which originate from proliferative expansions of individual cells and often possess altered cellular phenotypes such as resistance to therapies or pro-metastatic properties^3^. Deep sequencing has also revealed somatic mutations associated with autoimmunity and aging, including recurrent cancer-associated mutations even in histologically normal tissue^4–8^. However, the phenotypic impacts of these mutations remain poorly understood. Collectively, these observations motivate approaches that directly link somatic mutations to cellular phenotypes at single-cell resolution and that can be used to reconstruct somatic evolution.

Recent advances in single-cell sequencing have enabled the joint measurement of mutational genotypes and transcriptomic phenotypes within the same cell^9,10^, making it possible to define cellular lineages and subclones based on shared mutations and to characterize their functional cell states. Despite the promise of these single-cell multi-omic technologies, several shortcomings have limited their ability to resolve somatic evolution. In particular, existing methods for genotyping mutations from transcripts often amplify targeted loci after poly(A)-primed reverse transcription (RT), which does not scale beyond a few targeted sites (typically 1-3), restricts targets to mutations near transcript ends, and limits applicability to viable cells with intact transcripts^10^. Because genotype inference is highly sensitive even to rare errors, overcoming these limitations requires methods that are both scalable and highly accurate.

Here, we introduce Genotyping In Fixed Transcriptomes (GIFT) for single-cell multi-omic profiling. In GIFT, a unique reverse transcription reaction is used to fill targeted, mutation-bearing gaps between rationally designed single-stranded DNA (ssDNA), enabling simultaneous measurement of cell state and somatic mutations from the RNA of a single cell. We show that this approach overcomes limitations of prior RT-based methods, including the accurate capture of hundreds of loci in a single cell, and can be successfully applied to previously inaccessible formalin-fixed archival tissues. Profiling of 712,664 cells from 35 donors with myeloproliferative neoplasms (MPN) demonstrates that GIFT scales efficiently to hundreds of thousands of cells, recovers high-quality transcriptome profiles, and provides accurate and sensitive mutation inference irrespective of the position in the transcript. Across this cohort, we interrogate the phenotypic impacts of recurrent somatic mutations across hematopoietic differentiation. In individual patients, we use GIFT to reconstruct genetic lineages from multiple mutations and link relevant clones to diverse cell states. Together, single-cell multi-omic profiling via GIFT enables scalable, mutation-resolved inference of somatic evolution from heterogeneous human tissues and tumors.

## Results

Prior methods for somatic variant genotyping alongside single-cell transcriptomics have identified mutated cells in heterogeneous cell populations, but key technical limitations have precluded cohort-level profiling across diverse patient samples. Resolving clonal lineages—even at modest resolutions—requires capturing tens of mutations or more per patient, whereas existing approaches are limited to a few regions per patient. As a result, RNA-based genotyping has only provided a coarse view of clonal structure. Current approaches are also restricted to variants near the end of a transcript, limiting flexibility and scalability because mutations are often sparse and can occur anywhere along a transcript. These limitations are particularly prohibitive for resolving somatic mosaicism in normal tissue where relevant mutations are rare and distributed across many candidate loci^8^. These challenges are compounded by the incompatibility of existing technologies with FFPE samples, excluding a major set of clinically important human tissue.

The Single Cell Gene Expression Flex^11^ assay from 10x Genomics (‘Flex’ hereafter) enables gene expression profiling of fixed cells and nuclei, including from FFPE. In Flex, genes are targeted by pairs of probes that are hybridized and ligated to generate a barcoded molecule prior to sequencing (**Methods**), and whole transcriptome analysis (WTA) probe sets are commercially available for profiling human and mouse cells. Importantly, Flex probes can target any region of the coding sequence with minimal positional bias (**Supplementary Fig. 1a; Methods**). Given that established probe panels target ∼50,000 transcript sites, we reasoned that there would be substantial capacity to incorporate custom probes targeting hundreds or thousands of mutation-bearing regions, including candidate rare variants that may not be confidently defined from bulk sequencing. Furthermore, Flex supports scalable sample multiplexing for profiling large numbers of cells, a property that is essential for detecting rare clones in heterogeneous populations. These features suggested a genotyping strategy to directly read out native cellular genotypes using Flex-compatible probes, overcoming key limitations of existing methods. We implemented this strategy as an accurate, sensitive, and accessible platform for the genotyping of single cell transcriptomes at unprecedented scale, enabling mutation-resolved analysis of cellular phenotypes in primary and archival tissues.

### GIFT development and optimization

As Flex probe pairs hybridize to adjacent nucleotides on a transcript, we hypothesized that leaving a gap between probes could allow for a short segment of the native nucleic acid to be directly genotyped, even in fixed tissues. This strategy could be incorporated into the standard Flex workflow, including cell fixation, permeabilization, and probe hybridization, with two minor modifications—the addition of custom genotyping probes and inclusion of a gapfilling step to bridge the adjacent probe pairs—allowing both the transcriptome and genotyping probes to be ligated and processed for single-cell barcoding using droplet microfluidics (**Fig. 1a** and **Methods**). We term the integrated workflow, which combines whole-transcriptome gene expression profiling with targeted genotyping by ssDNA and gapfilling, Genotyping In Fixed Transcriptomes (GIFT).

**Figure 1.**
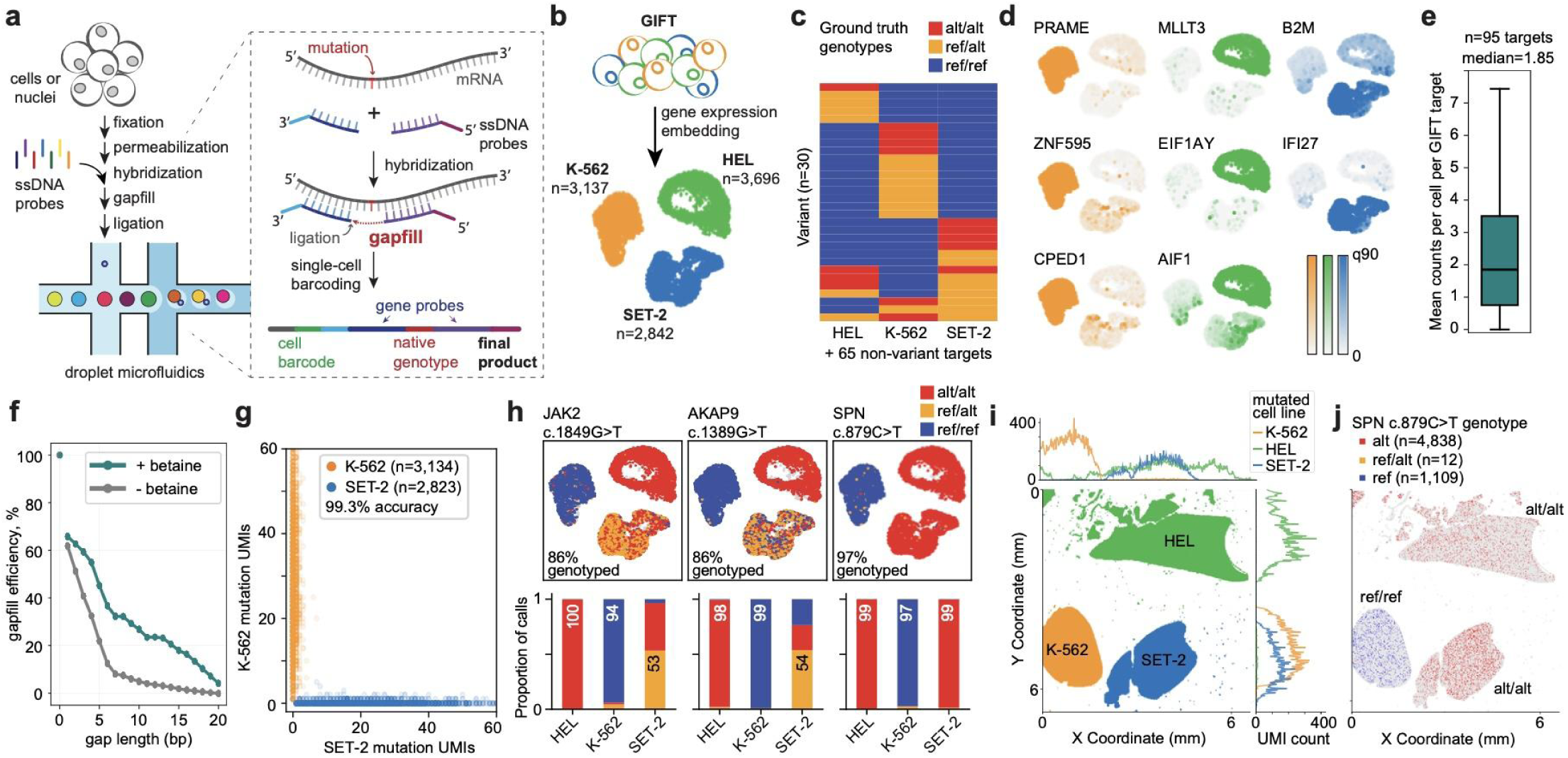
Gapfilling enables genotyping in addition to single-cell transcriptomic profiling at scale. **a,** The addition of a gapfilling reaction with ssDNA probes (dotted box) augments the conventional Flex workflow^11^ by capturing variants for single-cell genotyping at scale. **b-j**, GIFT profiling of a mixture of 3 leukemia cell lines with known genotypes. **b**, Uniform manifold approximation and projection (UMAP) showing that each cell type clusters distinctly by gene expression. **c**, Ground-truth genotypes for 30 variants in the GIFT panel. An additional 65 targets were not mutated in these lines (**Extended Data Table 1**). Ref, reference; alt, alternate allele. **d**, Expression of marker genes for K-562 (orange), HEL (green), and SET-2 (blue) cells. q90, 90^th^ percentile of normalized counts per cell. **e**, Counts per target that were captured per cell across the dataset. Line denotes median; box denotes interquartile range; whiskers extend to 1.5x interquartile range. **f**, Gapfill efficiency, the counts of a gapfilled probe pair relative to a 0-bp control probe on the same gene, depends on gap size and addition of betaine (locally estimated scatterplot smoothing from n = 301 probe pairs [15±9 per gap length]). **g**, Barnyard plot showing high single-cell purity and accuracy of GIFT genotyping (n = 5,957 cells). Cells are colored by transcriptomic clustering. **h**, *Top*: UMAP embeddings of three representative GIFT targets that vary across cell lines. *Bottom*: Genotyping accuracy for each variant. Numbers on stacked bars indicate the percent of genotyped cells with the correct call. **i**,**j**, Spatial genotyping with GIFT and Visium HD. **i**, Each 16 x 16 μm binned spatial feature is labeled by cell line as determined by gene expression, and histograms at top and right quantify GIFT counts for cell-line-specific mutations across X and Y coordinate axes. **j**, Spatial genotyping for a representative variant.

To establish GIFT, we sought a strand displacement deficient (SDD) polymerase capable of processing the missing nucleotides in the gap between adjacent probe pairs with no 3’→5’ or 5’→3’ exonuclease activity (**Supplementary Fig. 1b**), and we identified the DNA polymerase IV from the *Sulfolobus* genus as the best-fitting candidate (**Methods**). By reanalyzing existing human and murine Flex data, we developed an efficient computational workflow for custom ssDNA probe design to facilitate genotyping of arbitrary mutations, including preference for specific dinucleotide ligation junctions, balanced GC content, transcriptome-wide specificity, and gap size (**Supplementary Fig. 1c,d; Methods**). These custom probes enable creating a genotype-specific sub-library with minimal assay modifications (**Supplementary Fig. 1e; Methods**).

To assess the accuracy and sensitivity of GIFT, we designed a cell line mixing experiment combining the human HEL, K-562, and SET-2 cell lines and using a custom panel of gapfill-compatible probes to genotype 30 loci that were variable between cell lines and 65 additional sites with gap lengths spanning 2 to 10 nucleotides (**Fig. 1b,c; Extended Data Table 1**). Gene expression profiling yielded 9,675 cells with remarkably high-quality transcriptomic data (median 18,396 unique molecular identifiers (UMIs) per cell; median 5,976 genes/cell), including expression of cell line-specific marker genes (**Fig. 1d**). Importantly, inclusion of the gapfill enzymatic step did not meaningfully reduce gene expression (∼3.5% reduction; **Supplementary Fig. 2a,b**).

As part of assay optimization, we found that betaine, a common PCR additive^12^, increases GIFT yields per cell by 60% while maintaining high gene expression counts (**Supplementary Fig. 2c-e**). With betaine, we achieved a median of 1.85 counts per GIFT target per cell across 95 targets (**Fig. 1e**), enabling high confidence genotypes for most cells (**Supplementary Fig. 2f,g**). Variability in genotyping efficiency was largely explained by the target gene expression (**Supplementary Fig. 2h**). We next systematically defined the range of gap lengths that could be efficiently gapfilled using a multiplexed panel with gaps up to 20 bp (**Supplementary Fig. 2i; Methods**). While 98.5% of GIFT UMIs contained the intended gapfill size, gapfill efficiency decreased beyond 5 nucleotides, which was only partially mitigated by betaine (**Fig. 1f, Supplementary Fig. 2j**). From these data, our final probe design parameters incrementally penalize gaps longer than 4bp (**Methods**).

GIFT’s sensitivity was complemented by its accuracy, with 99.3% of counts reflecting the correct genotype in K-562 and SET-2 cell lines (**Fig. 1g**). Representative cell-line specific variants, including single nucleotide variants (SNVs) in the *JAK2*, *AKAP9*, and *SPN* genes, achieved 86-97% genotyping efficiency with high per cell accuracy (**Fig. 1h**). Moreover, heterozygous cells were correctly classified in >50% of genotyped cells for most variants, enabled by GIFT’s high per-cell depth that is necessary for characterizing heterozygous mutations and loss of heterozygosity (LOH) in patients^13,14^ (**Supplementary Fig. 2g**).

Similar to Flex, Visium HD spatial profiling utilizes short ssDNA probes to assay gene expression^15^. We therefore asked whether GIFT could be integrated into this workflow to enable spatially-resolved genotyping alongside transcriptomics. We adapted Visium HD for compatibility with GIFT and applied it to a spatially-restricted pattern of the same three cell lines (**Methods; Supplementary Fig. 3a,b**). Analyses of the GIFT library confirmed high accuracy (98.8-99.7%) from spatially resolved genotypes restricted to each respective cell line (**Fig. 1i,j**; **Supplementary Fig. 3c**). Consistent with differences in resolution between these modalities^15^, gene expression and GIFT complexity were lower in the spatial context than in dissociated single cells assayed by Flex (**Supplementary Fig. 3d,e**)^15^. Accordingly, we focused subsequent applications of GIFT on the dissociated single-cell workflow.

### Scalable and accurate genotyping with GIFT

While multiplexing of 95 loci already exceeds the genotyping scale of RT-based methods, the WTA panel contains ∼50,000 probe pairs to assay gene expression, suggesting that GIFT could scale to far more mutations to resolve clonal evolution through cumulative mutation acquisition. To test this, we created a GIFT genotyping panel targeting 611 loci and profiled four additional cell lines (Jurkat, HEK293T, HeLa, and MCF-7; **Fig. 2a**). Despite increasing the number of genotyping probes by nearly an order of magnitude, we retained high-quality transcriptional data, yielding 20,512 cells with gene expression counts per cell comparable to RT-based Genotyping of Transcriptomes (GoT)^10^ (**Fig. 2a**, **Supplementary Fig. 4a;** GIFT median for HeLa cells = 23,004 UMIs per cell). Gene expression enabled clear discrimination between cell populations alongside hundreds of genotyped targets per cell (median 164 GIFT targets captured per cell; **Fig. 2a-c**). In comparison to GoT, GIFT increased multiplexing ∼100-fold (**Fig. 2d**) without detectable transcript position bias (**Supplementary Fig. 4b**).

**Figure 2.**
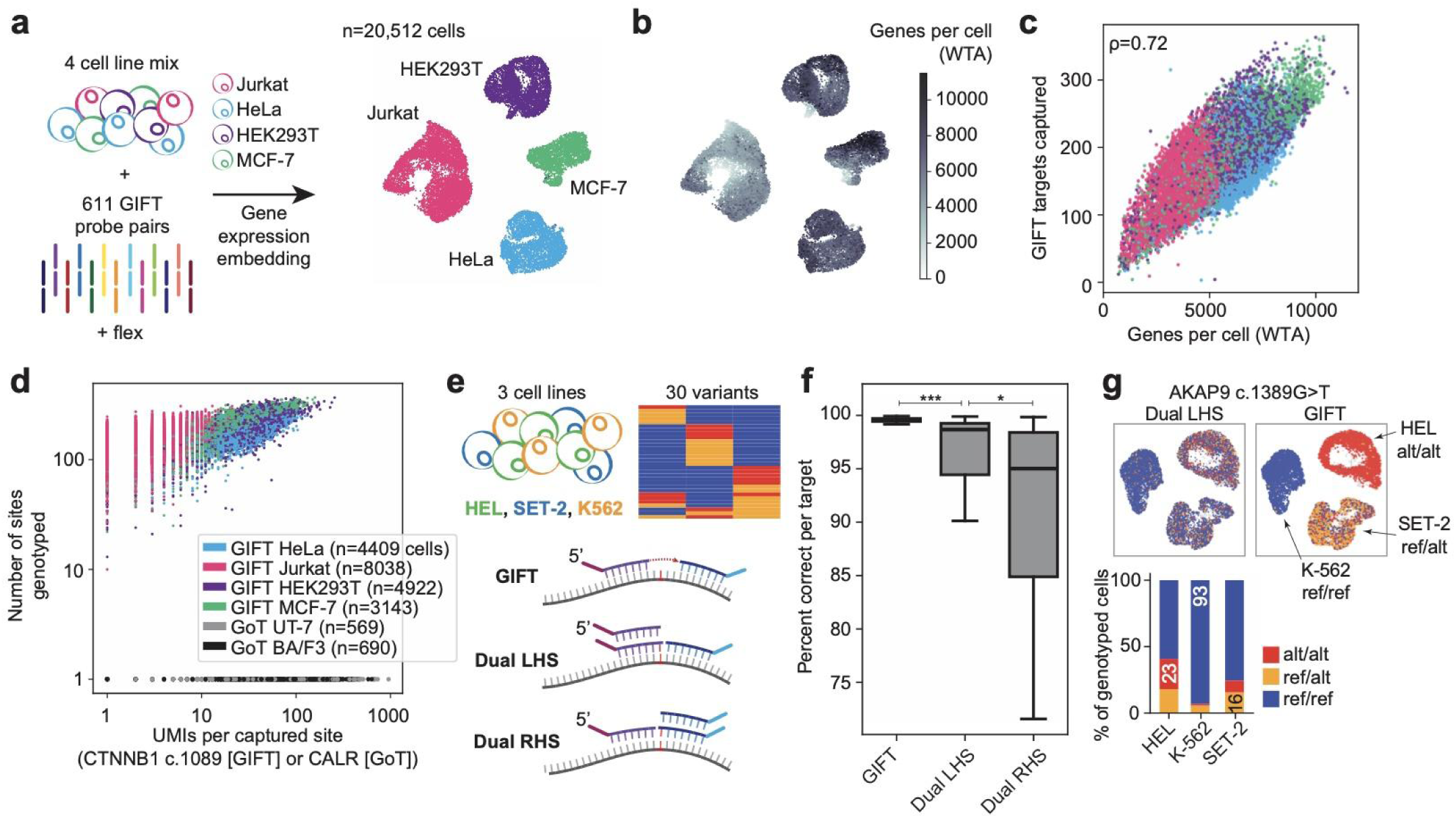
Scalable and accurate genotyping enabled by GIFT. **a–d**, GIFT profiling of 611 multiplexed targets in four mixed cell lines (n = 20,512 cells). **a**, UMAP embedding showing that each cell type clusters distinctly by gene expression. **b**, UMAP colored by number of detected genes per cell for gene expression assay (WTA, whole-transcriptome amplicon probe set). **c**, Relationship between number of GIFT loci genotyped per cell and number of genes detected per cell by WTA; ρ, Spearman correlation across all cells. **d**, Comparison of GIFT and GoT^10^ genotyping efficiencies. x-axis shows counts for the best-captured target in each dataset. **e–g**, Comparison of gapfilling and dual probes with WT and mutant alleles on the left-hand side (LHS) or right-hand side (RHS) probes in three mixed cell lines. **e**, Thirty variants were tested across the cell lines using the three methods. **f**, Percent of genotyping counts (UMIs) that are correct for each target by method (n = 30 targets per boxplot; *** p < 0.0005, * p < 0.05, one-sided Wilcoxon test). Heterozygous cells are excluded from accuracy calculations. Line denotes median; box denotes interquartile range; whiskers extend to 1.5x interquartile range. **g**, *Top*: UMAP embedding of a representative variant, colored by Dual LHS or GIFT genotypes (n = 11,187 genotyped cells by dual probes; n = 8,283 genotyped cells by GIFT). *Bottom*: Genotyping accuracy by dual probes, only considering cells with a genotype call. Numbers on stacked bars indicate the percent of cells with the correct call.

As an alternative to GIFT, the recently described GoT-Multi^16^ expands Flex for scalable probe-based genotyping using ssDNA probes complementary to both wildtype and mutant sequences of targeted loci (**Fig. 2e; Supplementary Fig. 4c**). Though this “dual probe” approach overcomes limitations of RT-based genotyping, including scalability and FFPE compatibility, we evaluated its accuracy relative to GIFT. As previously described^16^, dual probes improve genotyping accuracy when compared to single-allele targeting (**Supplementary Fig. 4d**). However, gapfilling substantially increases accuracy relative to dual probes with 90% of variants achieving >99% correct calls by GIFT versus 90.5% correct by dual probes (**Fig. 2f,g**).

Across replicates, GIFT is consistently more accurate than dual probes with minimal difference in genotyping counts per cell (**Supplementary Fig. 4e-h**). Taken together, these results show that GIFT combines the high specificity of RT-based genotyping with the scalability and flexibility of probe-based targeting.

### GIFT resolves somatic evolution in archival tissue

Although tissues preserved in formalin, including FFPE tissue blocks, represent one of the largest repositories of clinically annotated human material, they remain largely inaccessible to single-cell multi-omic analysis due to RNA degradation and crosslinking^17^. While Flex has enabled generation of high-quality gene expression profiles from archival FFPE samples, multi-omic extensions have been elusive, including the inference of somatic mutations from single-cell assays. We therefore asked whether GIFT could enable joint genotyping and transcriptomic profiling in archival tissue (**Fig. 3a**).

**Figure 3.**
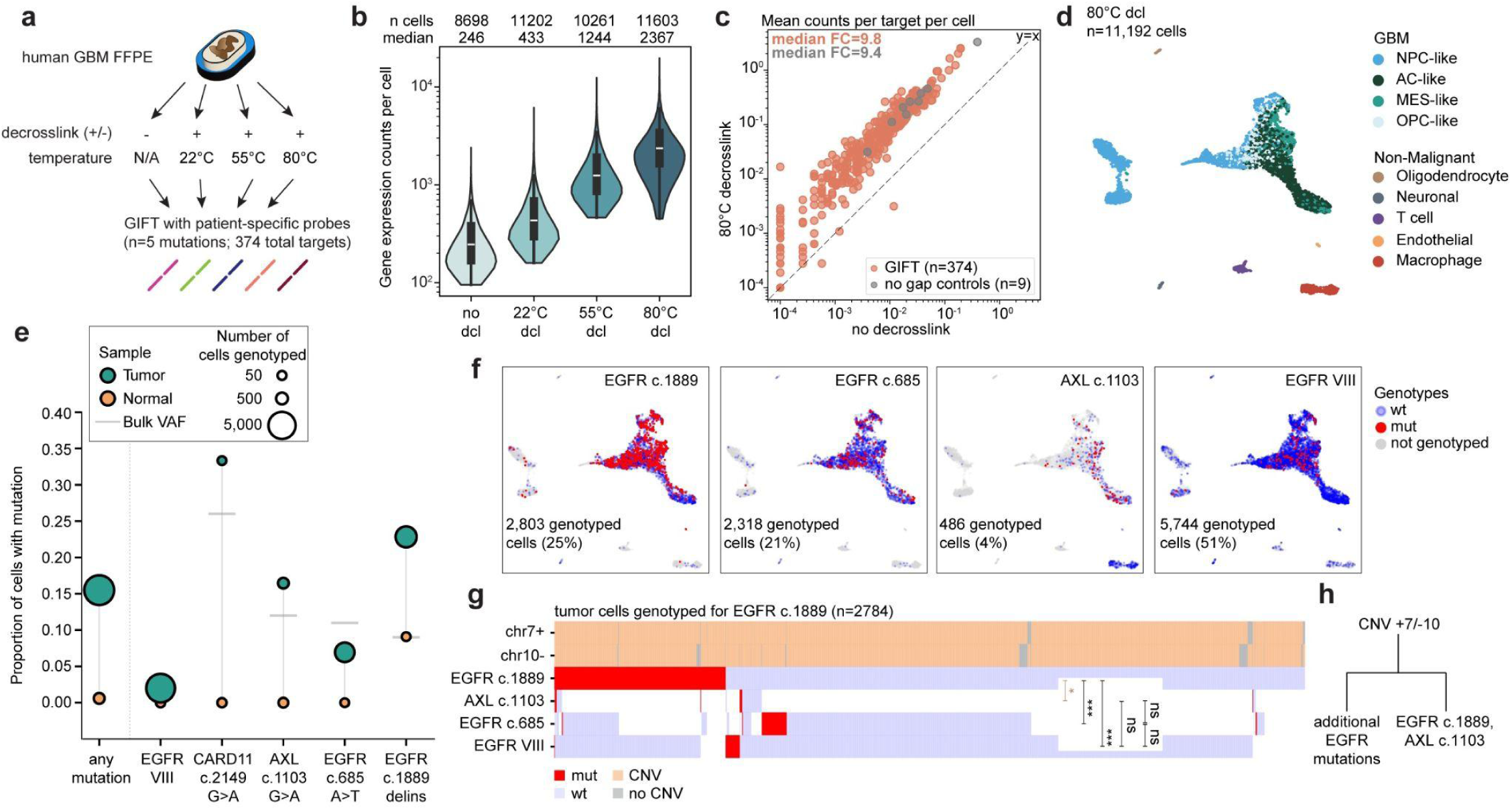
GIFT resolves somatic evolution in archival tissue. **a,** Human glioblastoma (GBM) FFPE tissue was subjected to nuclei extraction and multiple candidate decrosslinking conditions. **b,** Gene expression UMIs per cell for different decrosslinking (dcl) temperatures. Line denotes median; box denotes interquartile range; whiskers extend to 1.5x interquartile range. Conditions were compared in a single 4-plex experiment. **c**, Decrosslinking at 80°C increases GIFT counts per cell (*p* = 6.9*10^-62^, one-sided Wilcoxon test, n = 374). Gene-matched control probes containing the GIFT PCR handle but no gap are shown for comparison. FC, fold change. **d**, UMAP embedding of gene expression states from the 80°C decrosslinking library colored by cell type annotation (n = 11,192 cells). NPC, neural progenitor cell; AC, astrocyte; MES, mesenchymal; OPC, oligodendrocyte progenitor cell. **e**, (Pseudo)bulk variant allele frequencies (VAF) of mutations profiled from the cells in **d**. **f**, Genotype calls for major tumor-associated variants. The number of cells genotyped for each variant is noted with the percent of cells genotyped 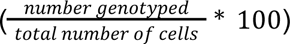. **g**, Mutational co-occurrence based on GIFT profiles of FFPE tumor cells, highlighting significantly co-occurring (*brown*) or mutually exclusive (*black*) mutations (Fisher’s exact test; *** p < 0.0005, * p < 0.05; number of cells as in **f**). **h,** Putative phylogenetic reconstruction of tumor evolution based on mutational co-occurrence.

We applied GIFT to FFPE-preserved human glioblastoma (GBM) tissue but initially recovered low gene expression counts per cell (median 246 counts per cell; **Fig. 3b, *left***). These low yields suggested that fixation-associated crosslinking limits probe accessibility. We therefore introduced a high-temperature decrosslinking step within the GIFT workflow to restore transcript accessibility for probe hybridization and ligation (**Fig. 3a; Methods**). This modification resulted in a ∼10-fold increase in gene expression recovery (median 2,367 counts per cell; **Fig. 3b**, *right*), indicating that this strategy substantially improves Flex gene expression profiling in FFPE. In parallel, GIFT genotyping yield increased by an order of magnitude (**Fig. 3c**), and gapfill efficiency was comparable to that observed in cell lines (**Supplementary Fig. 5a**), demonstrating that the underlying chemistry is effective in FFPE material. Importantly, the improvement in gene expression recovery was reproducible across two additional glioma patients (**Supplementary Fig. 5b-d**). Together, these results establish decrosslinking as a general strategy to restore transcript accessibility in FFPE samples, enabling robust probe-based gene expression and genotyping across archival tissue.

We recovered 11,192 filtered cells from the high-temperature decrosslinked GBM sample, which contained variable GBM subtypes and non-malignant cells, as defined by gene module scores and inferred copy number variants (CNVs)^18–22^ (**Fig. 3d; Supplementary Fig. 5e-h**). Accurate genotyping in FFPE samples requires distinguishing true mutation-derived signal from substantial technical noise, including artifacts from PCR-mediated template switching and errors introduced during gapfilling and sequencing.

Unlike gene expression measurements, which are relatively robust to occasional misassigned counts, genotype inference is highly sensitive to even rare errors, as incorrect allele assignments directly distort mutation prevalence, genotype–phenotype relationships, and clonal structure. We therefore developed a genotype-agnostic probabilistic framework to robustly recover genotype from noisy molecular observations that integrates evidence for each allele across four stages (**Supplementary Fig. 6a**; **Methods**). We first estimate UMI reliability and downweight molecules likely to reflect PCR-mediated template switching rather than true allelic signal (**Supplementary Fig. 6b-f**). We next restrict inference to a set of plausible alleles, pruning observed gapfill sequences to those consistent with biological signal rather than background noise, while enabling identification of sample-specific variants, including low-frequency events not captured by bulk sequencing. Each observed gapfill is assigned to the most likely allele, weighting calls by relative likelihoods, to recover signal even when gapfills do not exactly match the feature set (**Supplementary Fig. 6g-j**). Finally, we aggregate evidence across UMIs to infer per-cell genotype probabilities rather than discrete calls.

This framework is critical because false-positive mutation calls are especially damaging in the downstream analyses enabled by GIFT, particularly in challenging settings such as FFPE. By explicitly modeling UMI reliability, restricting inference to plausible candidate alleles, and integrating evidence probabilistically across molecules, our approach uses all available data while enabling thresholding at confidence levels tailored to downstream applications, a property that we exploit throughout our analyses.

For the GBM sample we defined a feature set including wildtype and mutant sequences for five targeted loci previously identified by bulk sequencing^23^, including three distinct *EGFR* mutations. Across confidence thresholds, false positive mutation calls in non-malignant cells decreased from 3.7% with the raw unthresholded output to 0.6% of cells at genotype probability above 60% (**Supplementary Fig. 5i**). This reduction in spurious calls is essential for reliable downstream analysis. Because the model scores homozygous and heterozygous genotypes, a 60% probability for a homozygous mutation typically reflects substantial support for mutation and a low probability that the cell is wildtype. In malignant cells, mutations were correspondingly enriched relative to non-malignant cells with 15% of genotyped cells including at least one mutation (**Fig. 3e,f**).

To assess clonal structure beyond bulk variant allele frequencies (VAFs), we quantified the co-occurrence of captured mutations to distinguish between sequential evolution in a single clone and parallel evolution in separate clones (**Fig. 3g**). We determined that an indel at *EGFR* c.1889 most likely co-occurs in the same clonal lineage as *AXL* c.1103G>A (**Fig. 3h**), thereby defining a distinct evolutionary branch from cells harboring the other two *EGFR* mutations. Whereas bulk data alone are compatible with 73 possible lineage trees, GIFT reduces this space to only six under standard clonal tracing assumptions (**Supplementary Fig. 5j**), demonstrating a substantial increase in resolving power.

Together, these results establish that GIFT enables mutation-resolved single-cell analysis in archival human tissue, a setting that has previously been inaccessible to multi-omic approaches. By combining improved sample preparation with probabilistic genotype inference, GIFT achieves the accuracy required to resolve clonal relationships directly from FFPE samples. This enables reconstruction of lineage structure and evolutionary trajectories within their native cellular context, directly linking genotype to lineage-resolved cellular states in a way not accessible to bulk sequencing.

### GIFT enables cohort-scale, high-resolution genotyping

The Flex kit enables a combinatorial pre-barcoding of transcripts, allowing single-cell profiling at cohort scale. To assess this, we sought to profile somatic evolution from patients with myeloproliferative neoplasms (MPNs), a group of blood cancers characterized by excessive proliferation of myeloid blood cells^24^. Unlike more aggressive malignancies, MPNs retain populations of canonical hematopoietic stem and progenitor cell states, providing a unique opportunity to assess phenotypic consequences of somatic mutations in a disease context with substantial relevance to normal and pre-malignant hematopoiesis, including clonal hematopoiesis of indeterminate potential^25^

The most common MPN driver mutation is *JAK2* V617F (c.1849G>T), which disrupts inhibitory control of the JAK2 pseudokinase domain^26^, followed in frequency by mutations in *CALR* and *MPL*, all of which cause constitutive JAK-STAT signaling^24^. Despite a shared initiating pathway, MPNs exhibit marked phenotypic and clinical heterogeneity, attributed in part to additional somatic mutations and to environmental influences^24^. Disentangling the phenotypic consequences of individual mutations within this complexity is a major challenge requiring high-resolution, multi-locus genotyping across many patients, which we reasoned could be tractable using GIFT.

To directly test this capability, we profiled a large cohort of 35 MPN patients and 2 healthy controls, generating whole-transcriptome and targeted genotyping profiles from 712,664 CD34^+^-enriched peripheral blood cells (median 14,330 cells per patient; range 1,205-55,604; **Fig. 4a**, **Extended Data Table 2**). Leveraging GIFT’s multiplexing capacity, we designed genotyping panels targeting nearly all known mutations in the cohort (n=67-71 targets per panel). GIFT recovered genotypes for up to 55 loci in a single cell (**Fig. 4b**), representing an order of magnitude increase in per-cell genotyping compared to prior studies with primary samples (3-8 loci per cell)^10,16,27^. In parallel, high-quality gene expression measurements enabled detailed identification of cell types, including CD34⁺ hematopoietic stem and progenitor cells (HSPCs) and differentiated myeloid, erythroid, and lymphoid populations^28^ (**Fig. 4c**; **Supplementary Fig. 7**). As expected, transcript abundance varied systematically across cell types (**Fig. 4d**), and GIFT capture similarly reflected gene expression differences, with the highest resolution in HSPCs (**Fig. 4e**).

**Figure 4.**
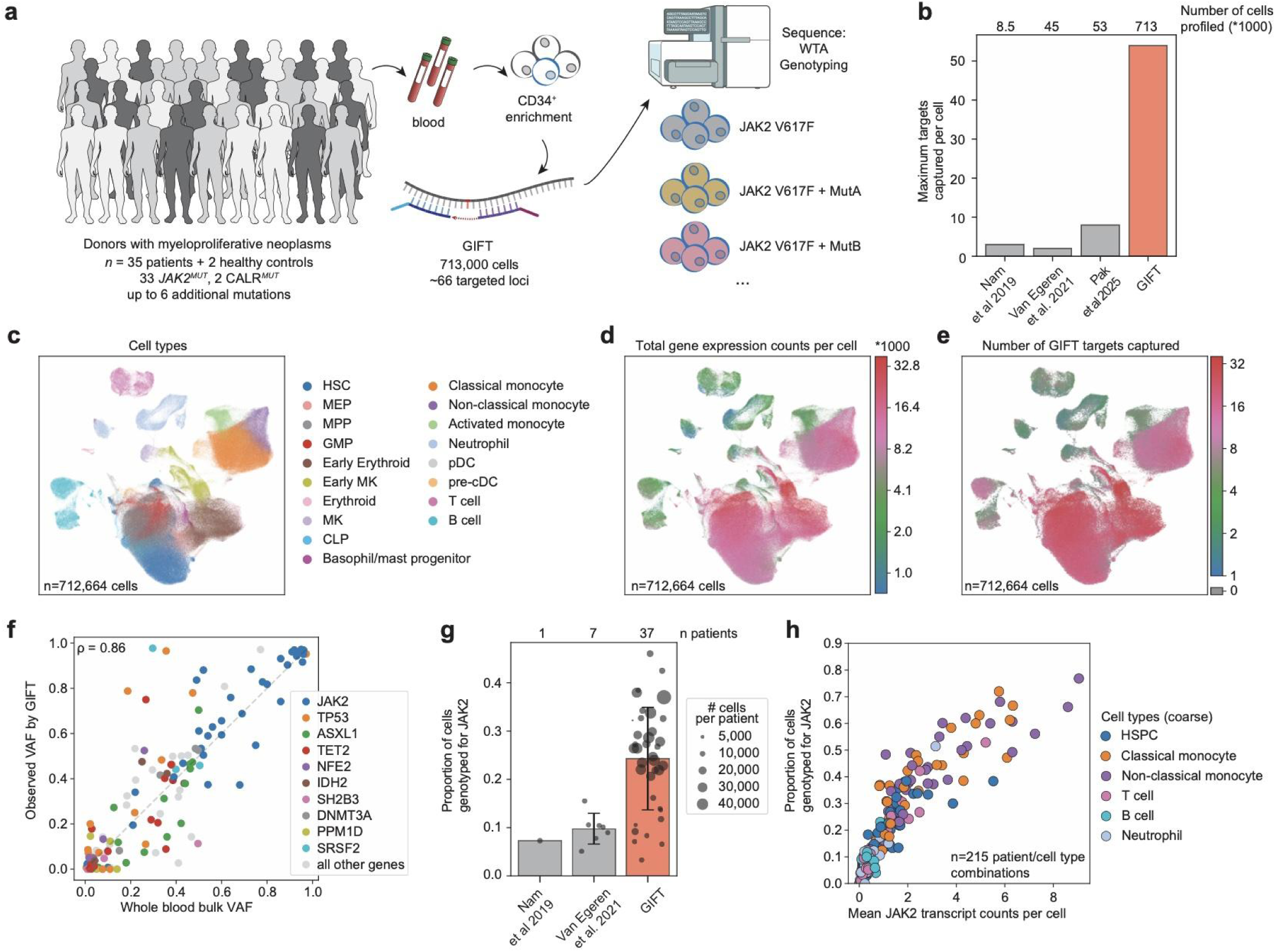
Cohort-scale profiling of myeloproliferative neoplasms with GIFT. **a**, Overview of MPN cohort. CD34^+^ peripheral blood cells from 35 patients and 2 healthy controls were profiled by GIFT. **b**, Number of targets and cells genotyped in primary samples with GIFT or three alternative approaches^16,27,54^. **c–e**, UMAP of patient-integrated scVI^59^ latent space, including all patients and controls, colored by cell type (**c**), total gene expression counts per cell (**d**) or number of captured GIFT targets (**e**). HSC, hematopoietic stem cell; MEP, megakaryocyte–erythroid progenitor; MPP, multipotent progenitor; GMP, granulocyte-monocyte progenitor; MK, megakaryocyte; CLP, common lymphoid progenitor; pDC, plasmacytoid dendritic cell; cDC, conventional dendritic cell. **f**, VAF measured by GIFT is highly correlated with bulk VAF in whole blood (ρ, Spearman correlation). Variants are shown only if expected from bulk (n = 138). Genes are labeled when more than 3 mutations in that gene are seen across patients. **g**, Proportion of cells (from all cell types) genotyped for JAK2 by GIFT (≥1 GIFT count) and two GoT-like approaches^27,54^. GIFT capture is significantly higher than alternatives^27^ (p = 0.0004, one-sided Mann-Whitney U). Error bars, s.d. across patients (*n* = 1 [Nam^10^], 7 [Van Egeren^27^], 37 GIFT—this study). HSPC, hematopoietic stem and progenitor cell. **h**, Proportion of cells genotyped for *JAK2* by GIFT is highly correlated with *JAK2* expression (Spearman’s *ρ* = 0.93, n=215 patient/cell type combinations).

Genotyping accuracy was validated by strong concordance between single-cell and bulk whole-blood VAFs across 138 expected variants (Spearman’s ρ = 0.86; **Fig. 4f**). Beyond detecting known mutations, GIFT enabled discovery of variants within targeted regions that were previously unannotated. For this cohort, we defined patient-specific feature sets by comparing the frequency of observed gapfill sequences in each sample to normal controls or to other samples, both of which proved effective for identifying novel candidate variants and filtering non-specific mutations (**Supplementary Fig. 8a-e; Methods**). Among two candidate discoveries (**Supplementary Fig. 8f**), an *NRAS* mutation (GIFT VAF = 0.003) was subsequently confirmed by bulk sequencing of this patient at a later timepoint (**Supplementary Fig. 8g**). In addition, GIFT uniquely enabled targeting of a *PPM1D* transcript region in which bulk sequencing found adjacent mutations (c.1636dup and c.1639del) in the same patient (**Supplementary Fig. 8h**). This flexibility of GIFT to target transcript regions rather than exact mutations is an important advantage over dual probes.

Our GIFT panels genotyped four sites that were profiled previously in MPNs by GoT, enabling direct comparison^10,27^. GIFT yields per cell were similar to GoT for variants near transcript ends, but GIFT dramatically improved capture of variants distal from transcript termini, including *JAK2* V617F (**Fig. 4g, Supplementary Fig. 8i-l**). Using GIFT, a mean of 24% of cells per patient were genotyped for *JAK2* V617F (≥1 GIFT count), corresponding to ∼3-fold increase over GoT-like methods^10,11^ (**Fig. 4g**). The proportion of cells genotyped for *JAK2* by GIFT closely tracked *JAK2* transcript abundance by patient and cell type, reinforcing that gene expression can be used to predict GIFT efficiency for new applications (Spearman’s ρ = 0.93; **Fig. 4h**). Consistent with prior reports, *JAK2* V617F cells showed preferential expression of the mutant allele relative to the wild-type transcript at the single-cell level^29^ (**Supplementary Fig. 8m**). Altogether, our results establish that GIFT enables scalable, accurate, and sensitive single-cell genotyping in primary samples while preserving high-quality gene expression profiles.

### Phenotypic impact of a recurrent JAK2 mutation

Understanding how somatic mutations alter cellular phenotypes in human disease requires measuring genotype and transcriptional state within the same cells at sufficient scale to disentangle mutation effects from patient-to-patient variability. In MPNs, recurrent driver mutations arise within a heterogeneous hematopoietic system, making it difficult to distinguish the direct effects of genotype from variation in cell type composition, disease stage, and patient-specific context^24^. A major limitation of prior approaches has been the inability to measure mutation status and transcriptional state at sufficient scale within the same individuals to resolve these effects. By enabling multiplexed genotyping across hundreds of thousands of single cells, GIFT provides a framework for directly comparing mutated and wildtype cells within shared cellular environments. We used this capability to investigate the phenotypic consequences of the *JAK2* V617F mutation in MPNs, applying a genotype-aware modeling framework to quantify both cell type abundance shifts and mutation-associated transcriptional programs.

In *JAK2*-mutant MPN, LOH by mitotic recombination at the *JAK2* locus commonly occurs as disease progresses^13,14^. To reduce ambiguity from heterozygous genotypes and isolate mutation-driven effects with high confidence, we leveraged the scale of our cohort to focus initially on 9 patients with predominantly homozygous *JAK2* mutation (**Supplementary Fig. 9a,b**). To quantify mutation-associated effects within this setting, we applied a deep generative modeling framework MrVI^30^ to construct latent representations of the data while explicitly accounting for sources of variation across patients. This approach allows us to distinguish variation attributable to mutation status from variation driven by patient-specific factors. In its standard formulation, mrVI learns a shared representation of cellular states across samples while accounting for structured differences between individuals, enabling separation of common biological programs from sample-specific variation^31^. Building on this framework, we incorporate genotype information directly into the model, treating mutation status as an explicit axis of variation, allowing us to construct representations of the data that either ignore or explicitly condition on mutation status. In practice, this yields two complementary views of the system: *u* that captures the baseline organization of cell states and *z* that reveals how those states are systematically altered by mutation. This enables direct comparison of mutated and wildtype cells within a shared representation, even in the presence of substantial inter-patient heterogeneity.

The genotype-unaware embedding, *u*, revealed differences in the abundance of *JAK2* V617F-mutant cells across cell types (**Fig. 5a,b**; **Methods**). Wildtype cells were enriched in lymphoid^32^ and pre-cDC cells, while mutated cells were enriched across multiple myeloid compartments, especially neutrophils (**Fig. 5c**). While neutrophils in *JAK2*-mutated MPNs have been reported to exhibit impaired apoptosis and contribute to high inflammation^33,34^, GIFT enables direct comparison of VAF across highly resolved cell types from the same patient. We also observed enrichment of *JAK2* mutation in early megakaryocyte (MK) cells and depletion in mature MK cells, consistent with a maturation bottleneck in mutant clones (**Fig. 5c**, **Supplementary Fig. 9c**)^35^.

**Figure 5.**
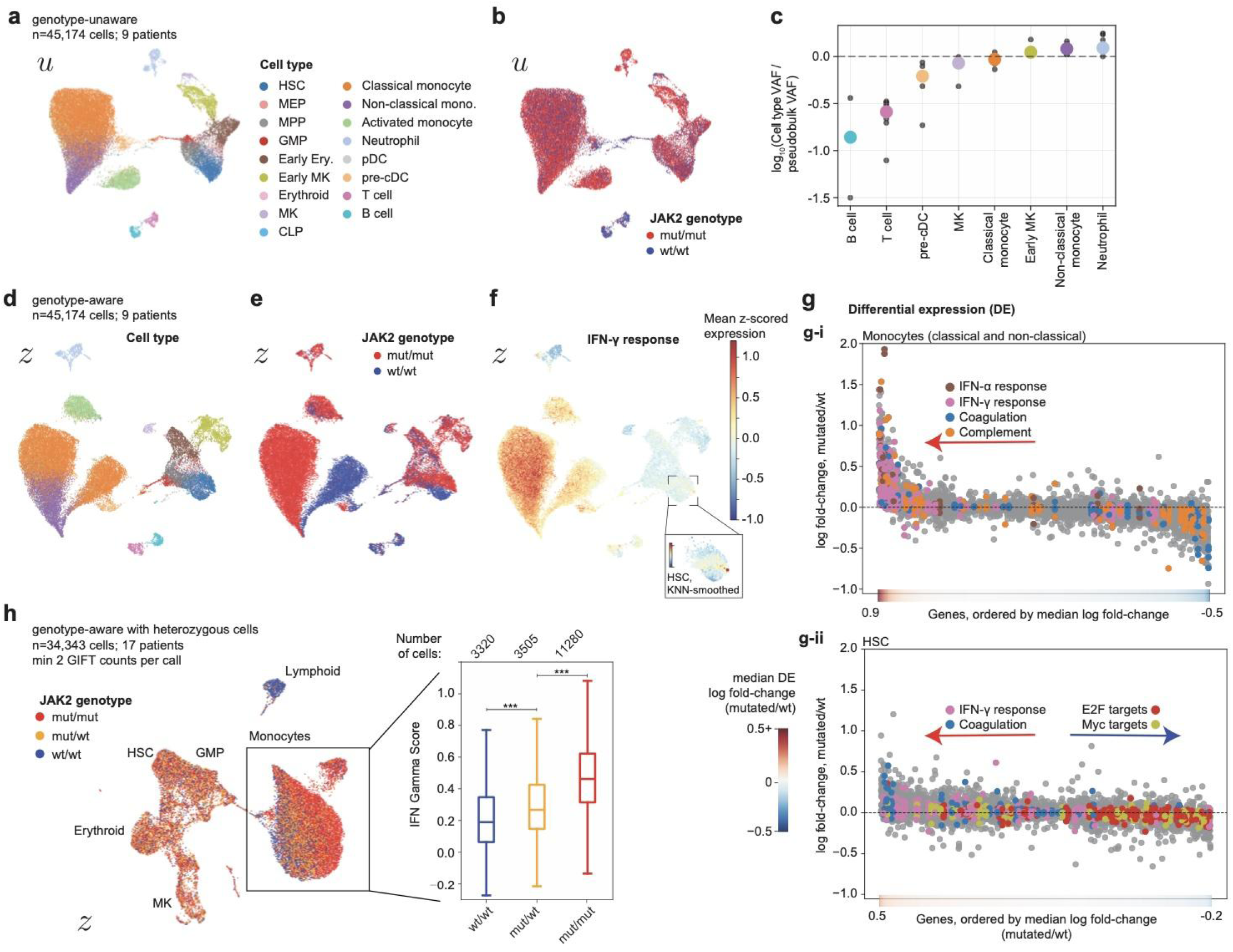
GIFT genotyping reveals impact of *JAK2* mutation in MPN patients. **a,b,** UMAP of genotype-unaware latent space (*u*) generated by MrVI^30^ for 9 MPN patients selected for having a low fraction of heterozygous cells, colored by cell type (**a**) or *JAK2* V617F genotype (**b**). Only non-heterozygous cells with ≥1 GIFT count are included (n = 45,174 cells). **c**, Differential abundance of *JAK2* V617F by cell type. *JAK2* V617F pseudobulk VAF by cell type is normalized to pseudobulk VAF across all cell types. Black dots show normalized VAFs for each patient (min. 20 genotyped cells), and larger colored dots show median across patients. Cell types are included if VAF significantly deviates from all-type VAF (p < 0.15, two-sided Wilcoxon test). **d–f**, UMAP of genotype-aware latent space (*z*), which isolates mutation-associated variation, generated by MrVI^30^ for the cells in **a**, colored by cell type (**d**), *JAK2* V617F genotype (**e**), or IFN-γ response gene score (**f**). Inset shows the KNN-weighted mean of gene scores for HSCs (k = 60) with a truncated color scale (ticks at −0.3 and 0.1). **g**, Differential gene expression in monocytes (*i*) and HSCs (*ii*). Each dot shows the log fold-change (LFC) of a given gene in an individual patient, ordered by median fold-change across patients (n ≥ 3 patients), and color bars (*bottom*) show median LFC. Labeled pathways^60^ are enriched in at least 3 patients (FDR < 0.2), and genes comprising each pathway are colored. **h**, UMAP of genotype-aware latent space (*z*) generated by MrVI^30^ including cells heterozygous for *JAK2* V617F. *Right:* IFN-γ response increases as dosage of *JAK2* V617F increases (one-sided Mann-Whitney U; p<10^-10^). Line denotes median; box denotes interquartile range; whiskers extend to 1.5x interquartile range.

We next examined mutation-associated transcriptional changes within specific cell types using the genotype-aware embedding, *z*, which separates mutation-driven effects from patient-specific variation within a shared representation (**Fig. 5d,e**). Because GIFT enables accurate, multi-locus genotyping across large numbers of cells within each patient, this framework allows direct comparison of mutated and wildtype cells within the same cellular contexts. Mutation-associated effects were observed across multiple hematopoietic compartments, most prominently in monocytes and hematopoietic stem cells (HSCs) (**Supplementary Fig. 10a**). Using MrVI counterfactual analysis, we quantified mutation-associated differential expression within monocytes and HSCs for each patient (**Methods**). Importantly, this method of differential expression analysis resolves differences across fine-grained cell-states without requiring discrete cell-level annotations. Across patients, we identified recurrently altered pathways despite substantial inter-patient heterogeneity, particularly for HSCs (**Fig. 5f,g; Supplementary Fig. 10b-d**). In both HSCs and monocytes, *JAK2*-mutant cells exhibit increased interferon-γ (IFN-γ) response (**Fig. 5f**), consistent with established links between constitutive JAK–STAT activation and inflammatory transcriptional programs^27,36–38^. Additionally, we observed reproducible upregulation of coagulation-associated genes in HSCs, which reflects megakaryocyte-biased hematopoiesis^36^ and notably persists in differentiated monocytes (**Fig. 5g**).

We observed depletion of E2F and MYC target gene programs in *JAK2*-mutated HSCs, suggesting relatively higher proliferation in wildtype HSCs (**Fig. 5g, Supplementary Fig. 10d**, **Supplementary Fig. 11a**). Notably, all patients included for HSC differential expression had myelofibrotic-stage disease with widespread LOH, and most were undergoing treatment, which can promote quiescence rather than proliferation in mutated cells^37,39^. Consistent with this interpretation, patients with non-fibrotic polycythemia vera (PV) and essential thrombocythemia (ET) exhibited higher HSC proliferation and more erythroid- and MK-biased hematopoiesis when compared to myelofibrotic disease, highlighting stage-associated differences in proliferation and output (**Supplementary Fig. 11b-d**).

We next extended the analysis to heterozygous cells across the full patient cohort to assess mutation dosage effects. Because accurate distinction between wildtype, heterozygous, and homozygous states requires sufficient genotyping depth, we restricted the analysis to cells with ≥2 *JAK2* GIFT counts per cell. Although this filtering removes 71% of genotyped cells, the scale of our dataset retains 34,343 cells for analysis. We then re-applied the genotype-aware MrVI modeling framework to this filtered dataset, generating a new genotype-aware representation (*z*) for these cells, analogous to the earlier analysis but now focused on resolving heterozygous and dosage-dependent effects (**Fig. 5h**, *left*; **Supplementary Fig. 10e-g**). Strikingly, we observed a gradient in genotypes within monocytes that was accompanied by a corresponding increase in IFN-γ response from wildtype to heterozygous to homozygous mutant cells (**Fig. 5h**, *right*; **Supplementary Fig. 10h**), expanding upon previously observed mutation-associated interferon signaling^38^. This direct observation of dosage-dependent phenotypes in primary human samples is enabled by the single-cell resolution, scalability, and sensitivity of GIFT, and would not be accessible using bulk or lower-resolution genotyping approaches.

### Lineage reconstruction in tumors

Somatic mutation profiling enables reconstruction of clonal evolution and identification of subclones linked to disease progression and drug resistance^3,40–42^, which we sought to resolve directly at single-cell resolution using GIFT. While bulk assays enable inference of clonal composition and phylogenies from allele frequencies^43^, single-cell profiling directly observes mutation co-occurrence within individual cells, enabling constrained reconstruction of lineage trees^44,45^. Beyond profiling mutations, GIFT further transforms clonal reconstruction by enabling direct attribution of cellular state and differentiation trajectories to specific clonal histories. These capabilities are particularly valuable in settings where rare malignant subclones emerge from broader populations of normal or less aggressive cell populations.

Clonal evolution is a central determinant of clinical heterogeneity in MPNs as this hierarchy influences prognosis and the risk of progression, including transformation to secondary acute myeloid leukemia (AML)^46–49^. In our cohort, we identified a patient undergoing progression from MPN to AML. UMAP embedding of cellular transcriptomes for this patient alone revealed distinct subpopulations corresponding to clones defined by four well-captured variants (**Fig. 6a**, **Supplementary Fig. 12a**). Differential expression analysis and comparison to longitudinal bulk genotyping enabled annotation of a leukemic blast-like compartment alongside an MPN compartment encompassing canonical HSPC populations (**Fig. 6b; Supplementary Fig. 13a-c**). Cells with heterozygous *CALR* mutation and/or homozygous *TP53* mutation were enriched in the leukemic blast but were still prevalent across MPN cells, highlighting phenotypic plasticity among cells with the same genotype (**Fig. 6c**).

**Figure 6.**
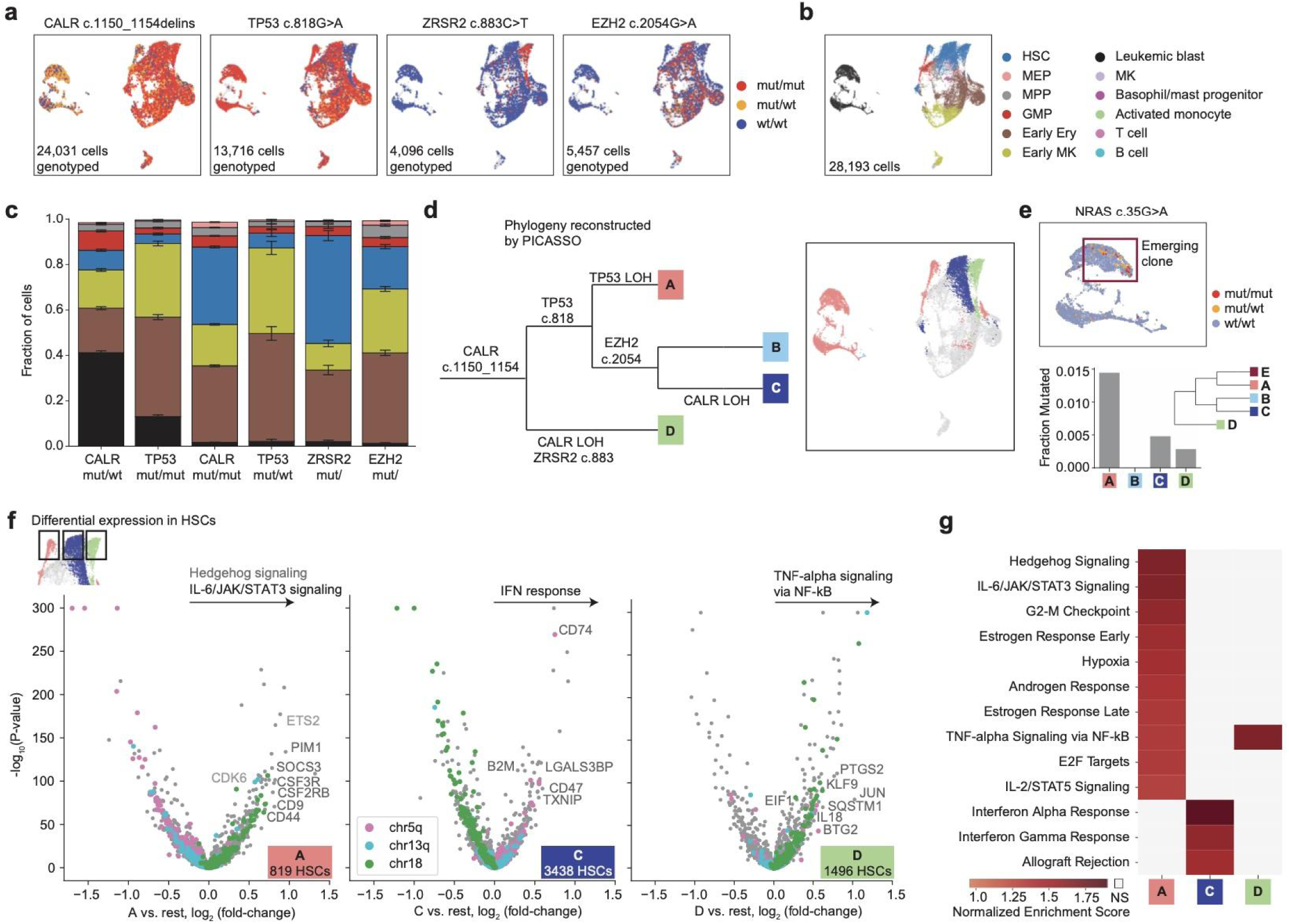
Lineage reconstruction in transforming MPN. **a,b**, UMAP of an individual MPN patient colored by GIFT-called genotypes for 4 different variants (**a**) or cell type (**b**). HSC, hematopoietic stem cell; MEP, megakaryocyte–erythroid progenitor; MPP, multipotent progenitor; GMP, granulocyte-monocyte progenitor; Early Ery, early erythroid; MK, megakaryocyte. **c**, Cell type fractions by cell genotype. Error bars, s.e.m. (n = 1-4,643 cells per condition). Colors correspond to the legend in **b**. *TP53* c.818G>A and *CALR* c.1150_1154delinsTGTC genotypes are classified as *mut*/*mut* or *mut*/*wt*, including only cells with p_genotype_ > 0.9. *ZRSR2* c.883C>T and *EZH2* c.2054G>A genotypes are classified as *mut*/ (*mut*/*mut* or *mut*/*wt*) with p_genotype_ > 0.6. **d**, *Left:* phylogeny inferred by PICASSO^44^. *Right*: UMAP colored by clone assignments. Grey populations could not be confidently assigned to a clone. **e**, *Top*: GIFT genotyping of *NRAS* c.35G>A. *Bottom*: *NRAS* c.35G>A mutation is enriched in clone A, which is the source of leukemic blast cells. We add E as an emerging clone defined by *NRAS* mutation. **f**, Differential expression^61^ in HSCs of clones A, C, and D relative to each other. The most significantly enriched pathways are shown for each clone, highlighting top pathway genes. Genes on chr5q, chr13q, and chr18 are colored to show effects of inferred CNVs. **g**, Normalized enrichment scores^62^ for significantly enriched pathways in HSCs of each clone (FDR <0.1). Grey boxes indicate no significant enrichment (FDR >0.1)

To assign cells to clones and infer a lineage tree, we used GIFT genotypes as input to PICASSO^44^, a probabilistic approach for phylogenetic reconstruction from single cells (**Fig. 6d, Supplementary Fig. 14a,b**). Accurate phylogenetic reconstruction critically depends on reliable genotype assignment, which is enabled by our probabilistic genotype inference framework. We excluded genotypes with confidence below 60% (**Supplementary Fig. 12b**) and only distinguished heterozygous from homozygous mutations for the *CALR* variant, which was sufficiently well-captured to genotype with >90% per-cell confidence (**Supplementary Fig. 14c,d**). These constraints on genotype confidence and allele representation prevent propagation of spurious genotype calls into phylogenetic structure. Without these constraints, the inferred phylogeny becomes uninterpretable and includes non-existent clones, such as a clone with both *TP53* c.818 and *ZRSR2* mutation arising from false positive genotype calls (**Supplementary Fig. 12c**).

Our clone identification was supported by expression-inferred CNVs that matched bulk karyotyping; clone A exhibited losses of chromosome 5q and chromosome 13q, and clone C showed loss of chromosome 18 (**Supplementary Fig. 15a-c**; **Extended Data Table 2**). The lineage tree includes a *CALR* heterozygous clone (clone A) and LOH yielding *CALR* homozygous clones (clones C-D). These LOH events are supported by high-confidence genotypes within each clone (**Supplementary Fig. 14e**) and by focal copy number loss near *CALR* (**Supplementary Fig. 15d**). For *TP53*, GIFT genotyping indicates LOH in clone A (**Supplementary Fig. 14f**), which is often observed in transformation to AML^50^. Absence of detectable chromosome-level loss is indicative of copy-neutral LOH^51^ (**Supplementary Fig. 15e**).

Remarkably, GIFT revealed incipient subclonal evolution preceding overt clinical progression. From our single-cell profiles, we identified a low-frequency *NRAS* c.35G>A mutation (enriched in clone A) not detected by whole-blood bulk sequencing (**Fig. 6e**). Longitudinal follow-up demonstrated expansion of this phylogenetic branch coincident with progression to post-MPN AML (**Supplementary Fig. 13c**), similar to established models in which acquisition of additional lesions, including RAS-pathway mutations, marks transition toward accelerated or blastic disease phases^50,52^. GIFT also enabled reliable detection of splice-site mutations with unknown functional consequences by targeting probes to pre-mRNA. Despite lower abundance relative to mature mRNA, we detected a splice-site mutation in *TP53* that was enriched in clone C. This was supported by reanalysis of bulk genotyping data and consistent with allelic imbalance of *TP53* c.818G>A in this clone (**Supplementary Fig. 13c,d**; **Supplementary Fig. 14g**).

Finally, coupling of high-resolution clonal ancestry with transcriptome-wide profiling allows for direct comparison of the expression patterns driving differentiation and proliferation while controlling for patient and microenvironment variation. In this single patient sample, we identified three HSC populations each corresponding to different clones (**Fig. 6f, Supplementary Fig. 13a**). Differential expression analysis revealed coherent transcriptional signatures of underlying CNVs alongside clone-specific gene programs (**Fig. 6f**). Strikingly, pathway analysis uncovered divergent biological programs across clones despite focusing exclusively on HSCs (**Fig. 6g**). The leukemic clone A was enriched for proliferative and inflammatory signatures, including hedgehog signaling and IL-6/JAK/STAT3 signaling, consistent with previous findings in post-MPN AML^52^. In contrast, the *EZH2*-mutated MPN clone C exhibited high expression of interferon response genes, and the *ZRSR2*-mutated MPN clone D exhibited activation of TNF-α signaling via NF-kB. Thus, mutation-resolved transcriptomes reveal programs that may direct clones along distinct differentiation and disease trajectories. Additional phylogenetic reconstructions across MPN patients further illustrate the robustness of our approach (**Supplementary Fig. 16**) and exemplify the novel biological directions enabled by GIFT. These analyses demonstrate that by integrating phylogeny and cellular phenotype at scale, GIFT reveals how distinct genetic clones drive divergent cellular programs and disease trajectories in primary human samples. Here, we define clones from up to six known mutations per patient, a scale that remains highly challenging for GoT even when all mutations are near the ends of transcripts. Looking forward, application of GIFT to genetically complex tumors with hundreds of variants will further enable phylogenetic reconstruction from somatic mutations at even higher, unprecedented resolution ^53^.

## Discussion

Coupling genotype to transcriptional state provides a rigorous framework to infer the impact of somatic mutations in clinical patient samples^54^. As the comparator (i.e., unmutated) population exists in the same individual and is profiled via the same single-cell library, many sources of variation can be effectively mitigated. Further, integrated profiles of cellular lineage and state can resolve functional tumor subclones that may evolve to accelerate disease onset, resist therapy, or metastasize^3^. These advantages collectively underscore the value of single-cell multi-omics that integrate genotyping and comprehensive cell state profiling. In recent years, several methods have emerged that pair genotypes to single-cell transcriptomic data^9,10,16,54,55^, but these approaches have been limited by minimal mutation multiplexing, biased position detection, incompatibility with formalin-preserved samples, low cell throughput, and/or complex assay design. Through a series of vignettes spanning primary and archived tissue samples, we demonstrate how the GIFT platform overcomes each of these limitations, offering a fully programmable platform for single-cell and spatial multi-omic profiling.

We demonstrate the successful detection of hundreds of genotyped sites in individual cells, a throughput previously not achieved by other transcriptome-based genotyping methods (to the best of our knowledge; **Fig. 2**). Additionally, GIFT utilizes rationally designed ssDNA probes that can efficiently hybridize across the transcript, mitigating 3’ detection bias from RT-based methods^10^. These ssDNA probes further enable genotyping heterogeneous input materials, including formalin-preserved tissues, such as FFPE blocks (**Fig. 3**). As our approach is built on top of a widely-used commercial platform with modest assay modifications, GIFT supports scalable sample multiplexing and practical adoption. In a cohort of MPN patients, GIFT enabled direct comparison of mutant and wild-type cells within shared cellular contexts (**Fig. 4**). These analyses revealed inflammatory transcriptional programs associated with JAK2-mutant cells and a dosage-dependent increase in interferon-response signatures in monocytes^38,56^, consistent with and expanding upon the known inflammatory signaling landscape of MPNs^38,57,58^ (**Fig. 5**). In a patient progressing from MPN to AML, GIFT resolved subclonal phylogeny together with clone-specific transcriptional programs and identified low-frequency mutations that later expanded during progression (**Fig. 6**)^50,52^. Together, distinct capabilities of GIFT enabled genotype-resolved profiling and biological insights that were previously inaccessible.

Looking forward, we anticipate that GIFT will aid in our continued understanding of somatic changes in healthy, pre-malignant, and diseased tissues. While deep DNA sequencing has discovered the recurrence of somatic mutations in both disease and ostensibly healthy tissues^1^, the impact of these mutations on cellular state has not been systematically defined. Pre-malignant and non-cancer settings present particular technical challenges because relevant mutations are distributed across many candidate loci and often only detectable by profiling large cell populations^8^. As GIFT readily scales to hundreds of targets and is built upon highly multiplexed cell barcoding, candidate variants can be comprehensively profiled simultaneously across large cell populations without extensive target-specific assay optimization. Together, the unique properties of GIFT position it as a formidable tool to define how somatic mutations shape cellular state and to resolve the accumulation of molecular changes in human tissues over a lifetime.

## ACKNOWLEDGEMENTS

We are grateful to the Pe’er Lab, Lareau Lab, and Single-cell Analytics Innovation Lab members for helpful discussions. We are grateful to T. Nawy for helpful feedback. We also thank I. Fiddes, J. Abousoud, T. Drennon, J. Sapida, N. Anaparthy, C. Kunihiro, and P. Smibert for additional contributions to this work. This work was supported by NIH grants P30CA008748 (C.A.L., R.C., D.P.), R00HG012579 (C.A.L.), U01AT012984 (C.A.L.), R37CA303960 (B.E.H., C.A.L.), R33CA302491 (C.A.L., R.C.), U54CA274492 (D.P.), P50CA257881 (D.P.), RM1HG011014 (D.P.), and Mark Foundation for Cancer Research Endeavor Award (D.P.). D.P. is a Howard Hughes Medical Institute investigator. SAIL is supported by the Alan and Sandra Gerry Metastasis and Tumor Ecosystems Center (GMTEC) and The Marie-Josée and Henry R. Kravis Cancer Ecosystems Project. S.B.B. is supported by a Junior Fellowship from the Simons Foundation. N.Ma. was supported by the ‘Ligue contre le Cancer’, the ‘Association pour la Recherche contre le Cancer’, and the ‘François Monahan Fellowship’. N.R. was supported as an HHMI-Fellow of the Damon Runyon Foundation for Cancer Research (DRG-2530-24). We acknowledge the Memorial Sloan Kettering Cancer Center Integrated Genomics Operation Core for sequencing services and the High-Performance Computing Group for computational resources. The funders had no role in study design, data collection and analysis, decision to publish or preparation of the manuscript.

## AUTHOR CONTRIBUTIONS

S.B.B., N.Ma., S.F., D.V., R.X., M.Mi., A.K., P.L., D.P., R.C., and C.A.L. conceived and designed the experiments. N.Ma., K.K., B.N., C.S., M.Ma., L.K., and M.T. performed experiments. S.F., N.Mo., E.B., D.V., L.B., R.X., M.Mi., S.W., N.S., L.D.-R., P.K., S.R., A.K., and P.L provided technical guidance and support for protocol development and optimization. K.K.H.Y., S.G., J.-J.K., and B.E.H. provided tissue samples and interpreted data. S.B.B. and A.V. developed software for probe design and data parsing with input from D.P. and C.A.L. S.B.B. led analyses with feedback from N.Ma., A.V., D.P., R.C., and C.A.L. with input from the other authors. S.B.B., N.Ma., A.V., D.P., R.C., and C.A.L. drafted the manuscript with input from other authors.

## COMPETING INTERESTS

S.F., N.Mo., E.B., D.V., L.B., R.X., M.Mi., S.W., N.S., L.D.-R., P.K., S.R., A.K., and P.L. are employees of 10x Genomics. D.P. is on the scientific advisory board of insitro. R.C. is a consultant for Sanavia Oncology and LevitasBio. C.A.L. is a scientific consultant to Cartography Biosciences. The remaining authors declare no conflicts of interest.

## METHODS

Our research complies with all relevant ethical and regulatory guidance. Recruitment of patients was approved under IDRCB 2023-A01414-41. Patient samples analyzed in this study were fully de-identified prior to transfer and analysis. The use of these specimens for the present work was reviewed and determined to be exempt from human subjects research by the Institutional Review Board of Memorial Sloan Kettering Cancer Center.

### Human cell lines

Human cell lines used in this work for mixing experiments included HEK293T, HEL, HeLa, Jurkat, K-562, Raji, MCF-7, SET-2. These lines were each procured from ATCC and monitored for mycoplasma contamination. Genotyping at ATCC confirmed the identity of all lines. Analyses of RNA expression in **Fig. 1-2** matched known gene expression and somatic mutation profiles, further confirming cell identity.

### MPN patient enrollment and sample collection

Peripheral blood samples were obtained from patients diagnosed with myeloproliferative neoplasms (MPNs and related disorders) enrolled in the Saint-Louis Hospital MPN cohort (Paris, France). All patients provided written informed consent for sample collection and research use in accordance with the Declaration of Helsinki and local regulatory requirements. The cohort is prospectively maintained, and peripheral blood samples are collected as part of routine clinical care. Peripheral blood samples were first processed by density gradient centrifugation using Ficoll to isolate mononuclear cells and were subsequently counted. CD34⁺ hematopoietic progenitor cells were isolated from peripheral blood mononuclear cells by immunomagnetic enrichment using the EasySep Human CD34 Positive Selection Kit (StemCell Technologies, catalog #17856), according to the manufacturer’s instructions. Following sorting, cells were counted and viability was assessed using a Luna FX7 automated cell counter (Life technologies), ensuring a post-sort viability greater than 80%. Purified CD34⁺ cells were cryopreserved in freezing medium and stored in liquid nitrogen for future research use. For this study, frozen aliquots were thawed, washed, and processed directly for single-cell profiling without additional in vitro culture.

### Targeted next-generation sequencing of whole blood samples

Targeted next-generation sequencing (NGS) was performed on DNA extracted from whole blood samples. A capture-based custom panel (Sophia Genetics) targeting 36 myeloid genes (ABL1, ASXL1, BRAF, CALR, CBL, CCND2, CEBPA, CSF3R, CUX1, DNMT3A, ETNK1, ETV6, EZH2, FLT3, HRAS, IDH1, IDH2, IKZF1, JAK2, KIT, KRAS, MPL, NFE2, NPM1, NRAS, PTPN11, RUNX1, SETBP1, SF3B1, SH2B3, SRSF2, TET2, TP53, U2AF1, WT1, ZRSR2) was used (Table S12). Libraries were prepared from 200 ng of genomic DNA isolated from whole blood cells (Qiagen) according to the manufacturer’s protocol. Sequencing was performed on a MiSeq instrument (Illumina). Bioinformatic analysis was conducted by Sophia Genetics (Switzerland) using the SOPHIA DDM software, and variants were reported at a threshold of 1%.

### GBM FFPE

FFPE samples were preprocessed on an S200+ S2 Singulator system. The sample was automatically processed in a NIC+ cartridge (S2 Genomics #100-215-389) by three 15 minute deparaffinization steps (CitriSolv, VWR), rehydrated by successive 1 mL washes of 100%, 100%, 70%, 50%, and 30% ethanol, followed by 2 washes of PBS. The sample was then spun at 1,000g for 3 min and resuspended in 0.5 mL Nuclei Isolation Reagent (NIR, S2 Genomics, #100-063-396) with 0.1 ul/uL RNase inhibitor (Protector, Millipore Sigma, #3335399001); all subsequent solutions had RNase inhibitor. The sample was dissociated to single nuclei in a second NIC+ cartridge with 2 mL of NIR for 10 min followed by a 2 mL wash with Nuclei Storage Reagent (NSR, S2 Genomics, #100-063-405). The single nuclei suspension was spun 500g for 5 min, resuspended in NSR, and counted.

### GIFT method and development

GIFT was developed to enable robust recovery of genotype information within the constraints of the Chromium GEM-X Flex v1 workflow (10x Genomics), using paired custom probes that define a variant-spanning gap subsequently copied by enzymatic gapfill (**Fig. 1a**) reaction. A complete end-to-end procedure for the GIFT GapFill workflow is available at protocols.io (https://www.protocols.io/view/gift-seq-hhz3b378p).

Method development focused on four technical axes that materially affected capture efficiency and library complexity: (a) probe design rules and junction context selection (see “GIFT probe design” and “Analysis of Flex WTA probes for custom design”), (b) permissible gap length, benchmarked across targets with matched 0 bp controls (**Fig. 1f**; **Supplementary Fig. 2i,j**), (c) enzyme selection to eliminate strand-displacement–driven boundary failure, and (d) nuclei handling optimizations, including post-sort decrosslinking to improve accessibility and downstream library performance (**Fig. 3b,c; Supplementary Fig. 5b,c**). The capture-improving technical components of the workflow are summarized below.

#### Sample-state optimization by freeze-thaw after fixation

During method development, we observed improved gapfill performance when the assay was performed on freeze-thawed fixed material following the fixation step of the Chromium GEM-X Flex v1 protocol (10x Genomics). This effect was observed across input types and was therefore incorporated as a general workflow optimization rather than a sample-type-specific modification. We hypothesize that freeze-thawing after fixation improves accessibility of probe-bound target molecules and facilitates more efficient enzymatic gapfill, thereby increasing recovery of genotype-derived molecules.

#### Identification of gapfill-compatible polymerase

In fixed tissues and cells, reverse transcriptase cannot process full RNA molecules, a requirement for most conventional scRNA-seq workflows. However, Flex utilizes a hybridization of short ssDNA probes that hybridize to target RNAs for quantification. We reasoned that we could infer a native genotype between subsequent ssDNA probes with a polymerase by genotyping over a short stretch (∼10 nucleotides) of an RNA molecule, even in fixed tissues. To identify a suitable enzyme that could achieve this gapfilling reaction, we focused on identifying an enzyme with the three key features: 1) no 3’→5 exonuclease activity;’ 2) no 5’→3’ exonuclease activity; and 3) no strand displacement activity (**Supplementary Fig. 1b**) Notably, the widely-used Moloney Murine Leukemia Virus (MMLV) RT is strand displacement competent^63^, which would displace the 3’ probe in a gapfill reaction. Thus, we surveyed potential enzymes commonly applied in synthesis reactions for activity in bridging pairs of ssDNA probes. We surveyed the set of candidate polymerases available from NEB (https://www.neb.com/en-us/tools-and-resources/selection-charts/dna-polymerase-selection-chart) and identified the *Sulfolobus* DNA polymerase IV as the only fitting candidate among polymerases for DNA manipulation. Thus, we developed and optimized our final gapfill reaction focusing on this enzymatic reaction.

#### Gapfill reaction optimization

The GIFT gapfill reaction was optimized to maximize recovery of probe-bounded genotype molecules. Probes were hybridized at 42°C for 16-24 hours. For some samples, hybridization started with a temperature ramp beginning at 60°C and decreasing by 1°C every 5 minutes until reaching 42°C; samples were then kept at 42°C for 16-24 hours. We did not observe a meaningful difference with or without this temperature ramp. Following post-hybridization washes, gapfill was performed on 100,000-120,000 cells per reaction using a master mix assembled on ice to ensure reproducible stoichiometry across multiplexed probe pools. The reaction combines ThermoPol buffer (New England Biolabs) and Flex internal amplification buffer (10x Genomics) to maintain enzymatic performance and downstream compatibility, dNTPs (New England Biolabs) to support complete fill across variable gap lengths (as benchmarked in **Supplementary Fig. 2j**), Protector RNase inhibitor (Sigma-Aldrich) to preserve nucleic acid integrity in fixed-cell/nuclei workflows, T4 DNA ligase (New England Biolabs) to stabilize the filled probe junction, and Sulfolobus DNA polymerase IV (New England Biolabs). Polymerase selection prioritized the absence of exonuclease and strand displacement activities (screened using the DNA Polymerase Selection Chart; New England Biolabs). The gapfill program (25 °C 10 min; 37 °C 20 min; 4 °C hold) separates an initial equilibration phase from an active extension/ligation phase, and reactions were diluted in post-hybridization resuspension buffer and washed once prior to GEM generation to minimize inhibitory carryover into downstream steps. Recovery of genotype-derived molecules was further improved by adding the TruSeq Read 2 reverse primer and performing two additional cycles during pre-amplification relative to standard Flex pre-amplification, followed by PCR enrichment of gapfill library and SPRIselect cleanup as specified in the protocols.io procedure.

#### Betaine as a gapfill yield enhancer

Betaine (Sigma-Aldrich), included in the gapfill master mix as specified in the protocols.io GapFill procedure, improved genotype molecule recovery (**Supplementary Fig. 2e,f**). Betaine reduces GC-dependent secondary structure formation and dampens melting-temperature disparities across probe-target duplexes, which decreases structure-driven polymerase pausing and incomplete fill events across heterogeneous loci. In practice, inclusion of betaine increased recoverable GIFT-seq library molecule counts while preserving compatibility with downstream Flex amplification and indexing.

#### Post-sort decrosslinking to improve nuclei-derived capture

For nuclei-derived inputs, we introduced a post-sort decrosslinking step that improved both WTA and GIFT library complexity and capture efficiency (**Fig. 3a, Supplementary Fig. 5b-d**; protocols.io: https://protocols.io/view/gift-seq-nuclei-preparation-hhjmb34k7). Following fixation/permeabilization and FACS enrichment, nuclei were washed in Sorting & Collection buffer (1× PBS, 0.2% BSA, RNase inhibitor), pelleted, and resuspended in freshly prepared decrosslinking buffer containing 150 nM 2-amino-5-methylbenzoic acid (162 nM in PBS), 0.5 M urea (8 M stock), and 0.5 µg mL^-1^ Proteinase K working solution (40 µg mL⁻¹; New England Biolabs), incubated at 80 °C for 30 min followed by 22 °C for 10 min, washed twice in PBS-T (1× PBS, 0.1% Tween-20), and resuspended in Quenching Buffer (10x Genomics) prior to starting Flex assay 1^st^ step probe hybridization. This treatment partially reverses fixation-associated protein–nucleic acid crosslinks while maintaining nuclear integrity, increasing probe accessibility, enzymes and improving downstream amplification efficiency, consistent with the measured gain in library recovery and quality improvement (**Fig. 3a, Supplementary Fig. 5b-d**).

### GIFT probe design

We developed a custom probe design pipeline for GIFT probes inspired by the 10x Genomics Technical Note CG000839 for standard, no-gap Flex probes. Probes were identified via score-based optimization of probe pairs, using stochastic dual-annealing in cases where explicit optimization was computationally prohibitive. Specifically, potential probe pairs (*l*, *r*) that capture a variant of interest within a gap of ≤10bp were scored. Probe pairs were excluded if the GC content of either side fell outside [0. 2, 0. 8]. A penalty score *S* was computed for each probe pair and minimized during optimization:

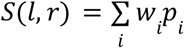

Where each penalty term *p* is defined as follows:

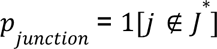 where 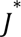 contains optimal junctions and *w_junction_* is tiered by suboptimal junction groups

*p_gaplengt_*_ℎ_ = 1[*l_gap_* ≤ 4] · |*l_gap_* − 4| where *l_gap_* is the length of the mutated sequence gap

*p_gc_* = |*GC_l_* − *GC_r_*| where *GC^l^*_/*r*_ is the proportion of G/C nucleotides in the LHS or RHS probe

*p_Tm_*_,*l*_ = *T_m_* (*l*) where *T_m_* is probe melting temperature

*p_Tm_*_,*r*_ = *T_m_* (*r*)

*p*_ℎ*omopolymer*_ = ℎ*max*(*l*) + ℎ*max*(*r*) where ℎ*max* is the maximum homopolymer length in each probe

*p_first_* = 1[*variant is first base of gap*]

*p_last_* = 1[*variant is last base of gap*]

*p_overlap_*_,*flex*_ = *n_overlap_* where *n_overlap_* is the number of bases in *l* and/or *r* that overlap with a probe in the WTA set

*p_overlap_*_,*GIFT*_ = 1[*any base overlaps wit*ℎ *anot*ℎ*er GIFT probe in t*ℎ*e panel*]

We initially determined weights (*w*) by mining the commercially available 10x Genomics WTA probe sets for Human and Mouse. We subsequently refined weights by post-hoc analysis of GIFT data. Specifically, we performed a regression of the average difference in genotyping and non-genotyping capture rate against the scoring parameters to compute optimal coefficients. A complete code base, including examples reproducing the probes designed in this study, is available in the **Code Availability** section.

### Analysis of Flex WTA probes for custom design

To inform GIFT probe design, we leveraged existing murine and human Flex WTA panels (10x Genomics), comprising around 50,000 probe pairs targeting protein-coding genes, typically with three probes per gene. We reasoned that multiple probes per gene provide replicate expression measurements that enable retrospective inference of probe-specific performance characteristics. Ensembl canonical transcript coordinates were obtained using biomaRt. Probe expression was quantified in mouse and human experiments across four tissues and analyzed in pseudobulk. We fit a linear mixed-effects model:

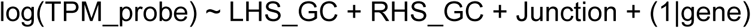

where junction was modeled as a categorical dinucleotide variable and gene identity was treated as a random effect. This framework allowed estimation of the contribution of probe attributes to residual expression variance. Consistent with commercial panel design, ‘NT’ junctions were strongly over-represented and associated with improved performance, and this constraint was incorporated into GIFT probe selection when feasible with fallbacks to secondary junctions guided by our analysis (**Supplementary Fig. 1c**). We also note that when ideal GC content is not available, probes can perform well between 20%-80% GC content, given that the LHS and RHS probes have similar GC contents.

### 10x Genomics Flex

10x Genomics Flex v1 was performed according to the manufacturer’s instructions with relevant modifications noted in **GIFT method and development** and **Supplementary Fig. 1e**. In addition to the human whole-transcriptome panel, a custom GIFT or dual probe panel was added at a concentration of 32 nM. For the GIFT versus dual probe comparisons, libraries were prepared identically except for addition of different custom probes. GIFT or dual probe libraries were sequenced alongside matched Flex WTA libraries on NovaSeq X (Illumina) or Aviti (Element Biosciences) using standard dual indexing. Read lengths were configured to fully span the probe-derived flanks and the filled gap sequence while maintaining the standard Flex barcode and UMI structure. Raw basecall files (.bcl/.bases) were converted to fastqs using bcl2fastq

(Illumina) or bases2fastq (Aviti) and processed with Cell Ranger v9 (10x Genomics), and WTA libraries were quantified against the human probe set v1.1.0 using default parameters.

### 10x Genomics Visium HD

Human HEL, K-562, and SET-2 cell lines were cultured and embedded into FFPE blocks. Tissue sections were cut at a thickness of 5 µm, spread in RNase-free water at 42°C, and mounted onto pre-prepared slides (Fisher Scientific #1255015). The slides were then air-dried at room temperature for 30 min and baked at 42 °C for 3 h. Subsequent experiments were carried out using dried slides stored for a few days at room temperature in a desiccator.

Briefly, tissue sections on glass slides were subjected to deparaffinization, H&E staining, and imaging following the Visium HD FFPE Tissue Preparation Handbook (CG000684, 10x Genomics). Probe hybridization, probe ligation, Visium HD slide preparation, probe release, extension, library construction, and sequencing were performed according to the Visium HD Spatial Gene Expression Reagent Kits User Guide (CG000685, 10x Genomics). After H&E staining, the tissue sections were destained and decrosslinked. The human whole-transcriptome probe panel was added to the tissue sections. In addition, a GIFT custom probe panel was added to the tissue sections at a final concentration of 24 nM per probe.

After hybridization, Probe Ligation Enzyme (PN-2000425, 10x Genomics) was added to establish connections between probe pairs hybridized to RNA, resulting in the formation of ligation products. After post-ligation wash, the Gapfill reaction was performed by adding Gapfill master mix to the tissue sections and incubating slides in a thermocycler at 25 °C for 10’ followed by 37°C for 20’ (For details: https://www.protocols.io/view/gift-seq-hhz3b378p). Following the Gapfill reaction, tissue sections were washed once with a 2× SSC buffer. Subsequent ligated probe release and capture within the 6.5 × 6.5 mm capture areas were performed using the Visium CytAssist. Treatment with RNase Enzyme and Perm Enzyme detached the single-stranded ligation products (WTA and GIFT probes) from the tissue and transferred them onto the Visium HD Slide for capture. These ligation products were then extended by adding Spatial Barcode, UMI, and partial Read1 primer. Subsequent elution and amplification of the ligation products prepared them for indexing through sample index PCR. Gapfill libraries were amplified separately using partial TruSeq Read1 and partial TruSeq Read2 primers. WTA and Gapfill libraries were cleaned and indexed separately using respectively Dual Index Plate TS Set A and Dual Index Plate TT Set A. Visium HD libraries were sequenced on the Element Aviti platform with dual indexing, and gene expression counts were estimated using the Visium v2 WTA probe set. SpaceRanger version 4.0.1 was used for analysis at 2 µm, 8 µm, and 16 µm resolution.

### Downstream analyses

#### Cell filtering, clustering, and visualization

Cells were filtered using WTA counts. All primary patient samples were first demultiplexed and then processed independently. Starting with the CellRanger filtered feature-barcode matrix, cells were further filtered by total counts (min = 1000, 500, or 100), genes (min = 500, 250, 100) and mitochondrial content (max=10%) for cell lines, MPN, and GBM samples, respectively. Genes were retained if detected in ≥55 cells. Raw counts were normalized to the median library size and log-transformed with base 2 and pseudocount of 1. We selected 4,000 highly variable genes (HVGs) using the highly_variable_genes function in Scanpy (v1.10.3, flavor=seurat v3)^64,65^. We performed principal component analysis (PCA) of log-normalized counts and retained 20 (cell line) or 30 (MPN, GBM) PCs. We clustered cells using PhenoGraph^66^ through Scanpy with the Leiden algorithm (k=200, resolution=1). We then identified cell doublets using Scrublet^67^ through Scanpy (expected doublet rate=0.06). We removed any barcodes predicted to be doublets as well as entire clusters with more than 60% predicted doublets. For MPN patient samples, we also removed cell clusters with mean mitochondrial content above 5%.

For cell lines (**Figs. 1-2**), GBM FFPE (**Fig. 3**), and individual MPN patients (**Fig. 6, Supplementary Fig. 16**), we re-processed the filtered matrix using the same steps as above (HVG selection, PCA, clustering).

Clustering parameters varied by sample (k=30 for **Fig. 6**, k=200 for all others; resolution=0.01, 1.5, 3, 1 for **Figs. 1-2**, **Fig. 3**, **Fig. 6**, **Supplementary Fig. 16**, respectively). We used the Scanpy neighbors function to compute a *k*-nearest neighbor graph (*k*=30, metric=Euclidean) and then the Scanpy UMAP function (min_dist=0.1-0.5) to generate a two-dimensional embedding. For GIFT and dual probes cell line mix UMAPs (**Fig. 1h**, **Fig. 2g**, **Supplementary Fig. 4h**), we processed both experiments together to generate one embedding. For the MPN cohort, we used the raw counts of the filtered matrix for data integration and visualization (see “MPN patient integration”).

#### GIFT Data Processing

The custom genotyping probes from the GIFT workflow are separately amplified and indexed, resulting in a separate library (.fastq.gz) that we parse with our *giftwrap* computational pipeline. At a high level, giftwrap parses the custom library to determine 1) the cell barcode, 2) the UMI, 3) the original designed probes, and4) the gapfill product with accompanying quality metrics. Giftwrap is implemented in a command-line compatible python library and distributed via pip. Efficient probe parsing is achieved via a pseudoalignment of reads through a prefix-trie index-based search. Cell barcodes are matched against a library-specific set of barcodes from the filtered feature-barcode matrix from either CellRanger and SpaceRanger for Flex and Visium, respectively. PCR duplication was resolved by grouping reads with matching cell barcodes and UMIs (within one edit distance). In rare instances where the same UMI matches to multiple probe pairs in the same cell, the most abundant is retained to reduce potential artifacts from PCR recombination. Finally, a consensus call of gapfilled sequences is determined across all PCR duplicates for each cell/UMI pair. Our toolkit additionally generates mapping summary statistics to enable quality control of GIFT libraries. Giftwrap provides an *scverse*-compatible Python interface^68^, utilities for generating a Seurat object^69^ from giftwrap processed data, and support for spatial genotypic data analysis through the *spatialdata* package^70^. Additional support for processing read-level gapfilled sequences is available as part of CellRanger version 9, which provides analogous parsing capabilities to giftwrap.

#### Assigning cell genotypes from GIFT counts

We employ a probabilistic framework to account for sources of error in GIFT data and assign probabilities to possible genotypes for each cell. This approach is intended to balance competing goals of sensitivity and accuracy. Rather than imposing initial UMI filters, we consider all available information in each dataset and generate probabilistic genotype classifications. This strategy enables application-specific downstream filtering and/or weighting of genotype classifications based on interpretable probability metrics. For example, our model computes probabilities of both heterozygous and homozygous mutation, which can be considered separately or combined to simply compute the probability of a mutation. For the phylogeny in **Figure 6d**, we filtered *CALR* genotypes to cells with >90% confidence, which enables discrimination of heterozygous from homozygous mutation, while for all other genotypes, we filtered to >60% confidence and collapsed heterozygous and homozygous mutation calls. This approach was essential for generating an accurate phylogeny from the available data (**Supplementary Fig. 12c** vs. **Supplementary Fig. 14a**).

As shown in **Supplementary Fig. 6a**, our probabilistic framework includes four steps. First, we compute the probability (*p_swap_*) that every detected UMI in a given dataset originated from template switching during PCR amplification. When template switching occurs, the cell barcode and UMI on one end of a contig form a chimera with the probe and/or gapfill of a different cell barcode and UMI, resulting in a sequenced output with no useful information^71^. Template switching occurs in any PCR-based library preparation, but it is particularly apparent in GIFT datasets because they are often low complexity and very deeply sequenced (saturation ∼95%). Additionally, gene expression profiling is more tolerant of occasional incorrect count assignments, which will not often meaningfully change the interpretation of a cell state, while genotype assignments are highly affected by any incorrect calls. As detailed below, we use the number of reads per UMI as a proxy to assign probabilities of template switching. An effective non-probabilistic strategy would be to impose read thresholds, as we do for processing cell line genotyping, but, to retain all information and maintain flexibility, we favor the probabilistic approach that we implement for all patient samples.

In the second step, we define a constrained feature set of possible mutations, which is essential to generate interpretable genotypes. As shown in **Supplementary Fig. 8a**, roughly the same number of gapfill counts from the MPN cohort match expected mutations as unexpected mutations. These unexpected gapfills span a large feature space of 202,493 observed sequences (vs. 138 expected mutations) and thus would add considerable noise if treated as true mutations (**Supplementary Fig. 8b**). The simplest solution here is to only consider expected mutations identified by bulk sequencing. However, we chose an empirical approach that retains only mutations with significant signal and enables discovery of new variants. In the MPN cohort, some expected mutations had much lower VAFs than bulk, which we could attribute in some cases (*TP53* c.733G>A, *TP53* c.659A>G) to differences in whole blood versus CD34^+^-enriched cells (**Supplementary Fig. 8f**, **Supplementary Fig. 12a**, **Extended Data Table 2**). More generally, this step to remedy discrepancies between GIFT and bulk VAFs is important in cases of different sample modalities and/or errors in clinical annotation. Additionally, we observed a few cases of expected mutations with high gapfill error rates. For example, *TET2* c.2908dupA was prevalent in healthy controls and across all samples and thus needed to be excluded from the feature set (**Supplementary Fig. 8c**). Finally, our empirical approach allows for variant discovery, which can reveal mutations missed by bulk sequencing or allow for GIFT profiling without bulk sequencing (**Supplementary Fig. 8f-g)**.

In the third step, we handle observed gapfill sequences that do not match sequences in the feature set. The simple solution here would be to remove these sequences, which we do for cell lines (**Figs. 1-2**). However, we noted that errant gapfills tend to be much more likely to originate from one included allele over others, as shown in the example in **Supplementary Fig. 6a,j**. Thus, we sought to instead retain these gapfill sequences and compute for each observed sequence the probability of originating from each candidate allele in the feature set. This step allows us to confidently assign the majority of unmatched sequences to the most likely allele while maintaining probabilistic outputs to account for uncertainty (**Supplementary Fig. 6g**).

In the final step, we integrate all UMIs for each cell to get cell-level genotype probabilities. Before this step, we have probabilities of the presence of an allele given an observed UMI. We use these values to determine the probability that a cell is wildtype, homozygous mutated, or heterozygous for each GIFT target.

Below we provide detailed computational methods for these steps.

##### 1. Computing probability of PCR-mediated template switching

We used a cell line dataset with confident ground-truth genotypes to assess the relationship between reads per UMI and genotype accuracy (**Supplementary Fig. 6b**). First, we computed the fraction of UMIs containing the correct gapfill as a function of reads per UMI (*n*). To convert these observed fractions into swap probabilities, *p_swap_*(*n*), we simulated permutations of the dataset. Specifically, we took a list of all observed UMIs in the cell line dataset and assigned each to the ground truth genotype for that cell type.

Then, we randomly permuted these assignments at rates *p_swap_* ∈ [0, 1] (**Supplementary Fig. 6c**). We recorded the resulting fraction correct and fit a linear regression, *p_swap_* = *f*(*fraction correct*) (**Supplementary Fig. 6d-e**). To then compute *p_swap_* for all *n* ∈ [*n_min_*, *n_max_*], we grouped observed cell line UMIs into non-overlapping bins of at least 2,000 observations, computed the mean read count (*n*), fraction correct, and corresponding *p^swap^* for each bin; we used local regression to interpolate between bins. To apply this mapping to a patient dataset, we sampled the cell line counts to match the depth of the patient data, using the knee points as normalization targets. We then computed *p_swap_* for each observed *n* in the patient dataset (**Supplementary Fig. 6f**).

##### 2. Defining a constrained variant feature set

We next defined a constrained feature set of gapfill sequences, *g*, for each probe pair, *p*, and sample, *s*. To identify gapfill sequences with significant signal, we considered three possible reference models:

1. “Control” model: Observed frequency of each gapfill sequence in healthy control samples. This model can be defined only when the relevant probe pair was included in the panel for controls, which was often but not always true for MPN targets (**Extended Data Tables 1-2**).
2. “Other” model: Observed frequency in other patient samples in the cohort. This model is more flexible because it does not require profiling healthy controls, and it is defined for nearly all targets in the MPN set. However, using other patients as a reference assumes that mutations should not be present in multiple patients, which would not be true for common driver mutations like *JAK2* V617F.
3. “Edit” model: Expected frequency based on the edit distance between the observed gapfill and the wildtype sequence. This model is the most flexible and is defined for any observed gapfill. As shown in **Supplementary Fig. 6h**, we compute explicitly the probability of sequential gapfill errors yielding the observed gapfill sequence from a wildtype template. Specifically, we iterate through possible gapfill errors, including truncations, insertions, or SNVs, using observed cell line frequencies as probabilities at each step (**Supplementary Fig. 6i-j**). Per-stage error probabilities are parameterized by error size (e.g., number of truncated bases or Hamming distance) rather than by specific nucleotide identity; this approximation is appropriate because the error rate distribution is dominated by a small number of common error configurations. The result of the edit distance calculation is the expected count (probability * number of probe UMIs) of the observed gapfill, which is the null expectation if all true alleles are wildtype.

For each model, we computed likelihoods of observed gapfills by a two-proportion Z-test comparing the observed gapfill count to the reference count under the model:

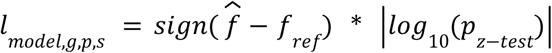

Where 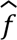 is the observed frequency and *f_ref_* is the frequency under the reference model.

To assess each model’s ability to discriminate signal from noise, we evaluated precision and recall for recovering expected mutations in the MPN cohort (**Supplementary Fig. 8d**). Our assumption here is that nearly all expected variants should be detected and that very few unexpected variants are true mutations. Models 1 and 2 (healthy control and other samples) both perform well with regions of very high precision (**Supplementary Fig. 8e**). Model 3 (edit distance) does not perform well for defining true variants and thus is only used in rare cases where neither controls nor other samples could be defined for that probe pair. Importantly, this highlights that profiling healthy controls and/or other samples with the same panel is necessary for robust mutation calling with GIFT.

To define patient feature sets, we applied a tiered selection strategy, prioritizing *l_control_*, then *l_ot_*_ℎ*er*_, then *l_edit_*, depending on availability. The constrained feature sets, *A_s_*_,*p*_, include all mutations expected from bulk with significant signal (*l_prioritized model_*_,*g*,*p*,*s*_ < −3). We then used a more conservative approach to add unexpected mutations to *A_s_*_,*p*_. First, we found the set of unexpected mutations with *l_control_*_,*g*,*p*,*s*_ < −25, number of observed gapfills >50, and frequency of gapfill >0.003. Then, we removed any mutations passing thresholds in more than one sample, reasoning that this was an indication of high noise for that mutation. Finally, we filter by *_ot_*_ℎ*er*,*g*,*p*,*s*_ < −50. For the MPN cohort, two unexpected variants were retained (**Supplementary Fig. 8f**) even with this very conservative filtering. For the GBM FFPE patient, we did not discover unexpected variants because we did not have a healthy control assayed with the same panel.

##### 3. Assigning observed gapfills to mutations in feature set

Next, for each UMI *i*, we computed the likelihood, *L*(*g_i_* |*a*), of observed gapfill *g_i_* under each candidate allele *a* ∈ *As*_,*p*_ using the edit distance model. Although this model is not reliable for distinguishing true variants from background (**Supplementary Fig. 8e**), it performs well for comparing relative likelihoods of the observed gapfill under a small set of (usually) 2 possible alleles. It is also defined in all cases, whereas it is not possible to use a healthy control to define *L*(*gi* |*mutation*) given that the mutation is not present in the control. To assign gapfills to their most likely allele, we normalize the likelihoods:

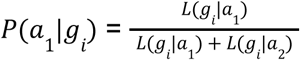

As shown in **Supplementary Fig. 6g**, most observed gapfills that do not match an allele in the feature set can be assigned by this method (P > 0.9) with high accuracy. For gapfills that do exactly match an allele in the feature set, this step also facilitates probabilistic discrimination between possible alleles.

##### 4. Integrating UMIs for cell-level genotypes

Finally, we integrate evidence across UMIs to obtain probabilistic cell-level genotypes. For each sample *s* and probe pair *p*, we score the probabilities of possible scenarios:

1. UMI originated from PCR swap and contains no relevant genotype information
2. UMI did not originate from PCR swap, and cell genotype *G* is:

a. Homozygous *a*_1_ (allele frequency α = 0)
b. Homozygous *a*_2_ (allele frequency α = 1)
c. Heterozygous (*a*_1_, *a*_2_) (allele frequency α = 0. 5)

We compute a score for each genotype *G* given each gapfill *gi* as follows:

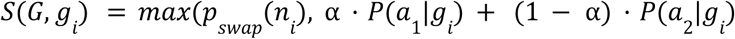

We use this maximum value heuristic because it is more conservative than a mixture model when *p_swap_* is high, though assignments are >99.9% consistent between these approaches. When *p_swap_* is high, it dominates *S* for all genotypes, resulting in uniform scores across genotypes.

We then approximate the per-cell genotype likelihood under *G* by taking the product across all UMIs from probe *p*:

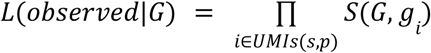

and compute posterior probabilities under a uniform prior:

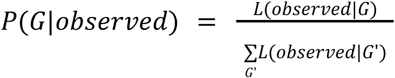

For discrete genotype assignments used throughout this work, each cell was assigned to *G* of maximum probability. Low confidence assignments were excluded (*P*(*G*|*observed*) < 0. 6). To count the number of cells genotyped or GIFT counts per cell, UMIs were included when *p_swap_* < 0. 1.

For cell line datasets (**Figs. 1,2**), we manually assigned per-library read thresholds to filter UMIs. We then set *p_swap_* = 0 for all retained UMIs. We did this instead of computing *p_swap_* because using cell line accuracy measures to assign *p_swap_* would introduce circularity in our benchmarking. We defined cell line feature sets using bulk genotypes from RNA-seq data downloaded from the Cancer Cell Line Encyclopedia^72^. In the interest of showing raw technical metrics from cell lines as clearly as possible, we did not use the edit distance model to assign observed cell line gapfills to alleles, even though this works well for cell lines (**Supplementary Fig. 6g**). Instead, we removed all UMIs with gapfills that did not match a wildtype or mutated sequence in the feature set. We computed cell-level genotypes as described but with *L*(*g^i^* ≠ *a_x_* |*a^x^*) = 0. 01 for GIFT and *L*(*g_i_* ≠ *a_x_* |*a_x_*) = 0. 1 for dual probes. Every cell with at least one retained UMI was assigned a genotype (no probability threshold because of pre-filtering).

#### MPN patient integration with scVI

For **Fig. 4**, filtered cells from all MPN patients were integrated using scVI^59^ (n = 712,664 cells). We identified the top 3,000 HVGs per sample using Scanpy (Seurat v3) and then retained HVGs (n = 4,021) that were shared between at least 10 of 37 samples. Raw counts were used as input to scVI using a 10-dimensional latent space and negative binomial gene likelihood model. Sample identity was included as a categorical covariate and mitochondrial read fraction as a continuous covariate. Using the scVI latent representation, we constructed a *k*-nearest neighbor graph (*k*=15) and then generated a two-dimensional embedding using the Scanpy UMAP function (min_dist=0.3).

#### MPN patient integration and genotype effect analysis with mrVI

MrVI is a deep generative model designed to evaluate effects of a target covariate while accounting for other nuisance covariates^30^. The model uses two levels of hierarchy to handle both types of covariate and learn two latent representations of the data. The *u* space is target-unaware and thus differences attributable to both target and nuisance variation are accounted for in this space. The *z* space is target-aware, so variation attributable to the target covariate is apparent, while nuisance variation is still integrated. In the original published implementation of MrVI, the authors demonstrate comparative analysis of patients with COVID-19 versus healthy patients. In this setting, the sample identifier is used as the target covariate because it is entirely nested in the target attribute of disease status; every sample has only one disease status. The model then enables differential abundance and differential expression analysis through counterfactual representations of disease status.

In our study of the effects of *JAK2* mutation in MPN patients (**Fig. 5**), MrVI provides a useful framework for handling genotype- and sample-level variation after important modifications to the typical setup. Our target attribute is *JAK2* genotype, which varies across the cells of the same patient, unlike disease status in the example above. Thus, we instead model sample identity as a nuisance factor and use genotype directly as target covariate, allowing MrVI to model genotype differences without effects of sample identity. We can then use counterfactual analysis to compare gene expression of the same cell in *u* space when mapped to *z* space as a wildtype or mutated cell. This method of differential expression analysis accounts for cell-level differences (e.g. cell type) without requiring discrete cell-level annotations.

In our first implementation of MrVI (**Fig. 5a-g**), we focused on the effect of *JAK2* mutation in wildtype versus homozygous mutated cells. To do this, we identified patients that had undergone LOH and thus had a low frequency of heterozygous cells. This constraint allows us to consider all genotyped cells, even if they only have 1 genotype count, and to remove noise from heterozygous genotyping (**Supplementary Fig. 9b**, MPN25 and MPN3). Specifically, samples were included if they met the following criteria for *JAK2* V617F, assessed among non-T and non-B cells with ≥ 2 genotype counts: (1) heterozygous fraction < 20%, (2) mutated cell fraction > 5%, and (3) wild-type cell fraction > 5%. After defining this patient set, we then included all cells in these samples with ≥ 1 genotype count for *JAK2* V617F unless genotyped as heterozygous, resulting in 45,174 included cells. Patients with fewer than 100 wt/wt or mut/mut cells were then removed, leaving 9 samples (**Supplementary Fig. 9a**). We identified the top 3,000 HVGs per sample using Scanpy (Seurat v3) and then retained HVGs shared between at least 3 samples (n=3,990 genes).

We also implemented MrVI to focus on dosage-related expression changes from heterozygous to homozygous mutated cells (**Fig. 5h**). In this implementation, genotype resolution is limited in HSPCs, evidenced by mixed genotypes across the gene expression space, which highlights why limiting the patient set in our first analysis was necessary. However, in monocytes, which have the highest *JAK2* genotyping counts and the greatest gene expression response to *JAK2* mutation, we observe a gradient of expression changes from wildtype to heterozygous to homozygous mutated cells (**Fig. 5h**). To implement MrVI with heterozygous cells, we carried out the same steps as the first implementation but with a few modifications. First, only cells with ≥ 2 genotype counts for *JAK2* V617F were included, resulting in 55,524 cells. Then, samples were retained if they included at least 100 genotyped cells and if wildtype and mutated cell fractions were >5%, resulting in 17 samples containing a total of 34,343 cells. HVGs were retained if shared between at least 5 (of 17 total) samples (n=4,166 genes).

For both implementations, raw counts from the filtered, concatenated dataset were used as input to MrVI with a 10-dimensional *u* and 30-dimensional *z* latent space. As described, genotype was used as the sample key and sample as the batch key (nuisance factor). Two latent representations were extracted from the trained MrVI model, a genotype-unaware representation, *u*, and a genotype-aware representation, *z*. For each representation, we constructed a *k*-nearest neighbor graph (*k*=15) and computed UMAP embeddings (min_dist=0.3).

#### Cell type assignment

To annotate cell types for the GBM patient (**Fig. 3**), we first distinguished malignant from non-malignant cells.

We used infercnvpy, a python implementation of inferCNV^21^. Log-normalized counts for all genes (pre-filtering) across filtered cells were used as input. CNV inference was performed with a sliding window of 100 genes and step size of 1, using a cell cluster expressing T cell genes^18^ as a non-malignant reference.

The resulting CNV matrix was used as input for PCA and clustering by PhenoGraph. Clusters were assigned as malignant or non-malignant based on abundance of the pre-defined reference cells and relative CNV burdens. To define +7/-10 cells in **Fig. 3g**, per-cell mean CNV scores were computed across chromosome 7 and chromosome 10, and thresholds were set by visual inspection (**Supplementary Fig. 5f**). Cell types were assigned by computing gene scores as the mean z-scored log-normalized expression across previously defined gene modules^18–20^. Non-malignant cell types were assigned at the cluster level, while malignant cells were individually assigned to the highest scoring GBM transcriptional state.

Cell identities in the MPN cohort (**Fig. 4–6**) were assigned using a supervised workflow combining manual annotation with automated classification, guided by canonical lineage-marker expression and comparison to reference atlases of human hematopoiesis^28,73^. We first manually annotated clusters from two reference patients spanning the major hematopoietic compartments. Early hematopoietic stem and progenitor states were defined by enrichment of stemness programs (e.g. *CD34, HLF*) and lineage-priming signatures (e.g. *MPO*, *ITGA2B*, *GATA2*) resolving myeloid, erythroid–megakaryocytic and lymphoid trajectories (**Supplementary Fig. 7a,b**). Mature immune and differentiated populations were annotated using established lineage-defining markers across lymphoid, myeloid and dendritic compartments (**Supplementary Fig. 7c**). We deliberately relied on combinations of lineage-consistent genes rather than any single transcript to minimize overannotation of transitional states, particularly within the progenitor and dendritic-cell compartments. Marker sets used for annotation are summarized in **Extended Data Table 3**.

These curated cells were used to train a custom CellTypist classifier^67^, subsampling up to 500 cells per cell type. The trained model was then applied independently to each patient to generate per-cell predictions, which were subsequently reviewed and refined on the basis of canonical marker-gene expression patterns.

Malignant blast populations (**Fig. 6b**) were annotated separately from normal hematopoietic cell states based on their transcriptional profiles and concordance with clonal structure inferred from bulk phylogeny (**Supplementary Fig. 13a-c**), consistent with previously described leukemic blast transcriptional programs in single-cell studies of myeloid malignancies^74,75^.

#### Genotyping accuracy

To compute the accuracy shown in **Fig. 1g**, we considered genotyped targets that were mutated (heterozygous or homozygous by bulk reference) in either SET-2 or K-562 cells, excluding any mutations shared by both cell lines, and computed accuracy across all targets:

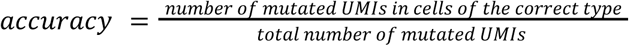

To compute the percent correct shown in **Fig. 2f** and **Supplementary Fig. 4d,e**, we considered genotyped targets that were mutated (heterozygous or homozygous by bulk reference) in at least one cell type and computed the percent correct for each target:

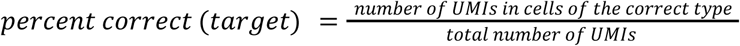

Because of inconsistencies in bulk genotyping, we excluded *HTT* c.8157G>A from both metrics above.

#### Differential abundance analysis

For differential abundance analysis shown in **Fig. 5c**, we considered each patient sample *s* in the integrated dataset (**Fig. 5a-b**) and cell type *k* with at least 20 genotyped cells in that sample. Thus, we considered a different number of samples, ranging from 1-9, when considering each cell type; for example, all 9 patients included in **Fig. 5a-b** had ≥ 20 Non-classical and Classical monocytes, but only 1 patient had ≥ 20 Erythroid cells.

We then computed *f_s_*_,*k*_, the proportion of mutated cells in cell type *k* in sample *s*, and *f_s_*, the proportion of mutated cells across all cell types in *s*. We filtered for cell types in which *f_s_*_,*k*_ differed significantly from *f_s_*, indicating significant enrichment or depletion of *JAK2* mutation in that cell type. Specifically, we used a two-sided Wilcoxon test (matched by sample) and included cell types with p < 0.15 (n = 8 cell types, each covered by 4-9 samples).

For each included cell type, we quantified enrichment of the *JAK2* V617F mutation as log fold-change (LFC) relative to the sample-level pseudobulk mutant allele frequency:

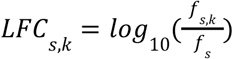

Median LFC values by cell type are shown by colored dots in **Fig. 5c** with each sample shown by grey dots.

#### Differential expression analysis

We used MrVI^30^ to compute per-cell counterfactual differential expression between *JAK2* V617F mutant and wildtype cells (**Fig. 5g**). Log fold changes (LFC, base 2) are computed by mapping each cell’s genotype-unaware representation (*u*) to alternative genotype-specific (*z*) states and comparing the decoded gene expression. For differential expression analysis, we re-implemented MrVI as described above for each patient individually (only for the 9 patients included in **Fig. 5a-f**; 2,000 HVGs retained per patient). We did this instead of using the patient-integrated MrVI model because the number of cells varies substantially across patients (n=387-20,146 total cells per patient) with even more variability when broken down by cell type. Implementing the model separately for each patient allowed us to consider all cells from every patient while being very careful to avoid a few patients dominating the *u* to *z* mapping.

We focused our analysis on HSCs and monocytes (classical and non-classical) and required at least 30 cells per genotype per cell type for a patient to be included (n = 6 patients for monocytes, n = 5 patients for HSCs). Mean cell type LFCs (for each patient) were used as input to the prerank function of GSEApy^62^ with MSigDB Hallmark gene sets^60^. Pathways with FDR (q-value) < 0.2 were considered significant. We used this relatively lenient FDR because we subsequently focused on pathways significant in at least 3 patients, making the convergent set substantially more stringent (**Fig. 5g**). The full set of pathways significant for at least 1 patient is shown in **Supplementary Fig. 10c,d**.

To compute differential gene expression between phylogenetic clones, we first restricted each comparison to a single cell type (HSCs in **Fig. 6f** and **Supplementary Fig. 16j**; classical monocytes in **Supplementary Fig. 16d**) in order to focus the analysis on clone rather than cell type differences. For the three clones in **Fig. 6f**, we compared each clone to both other clones and focused only on upregulated genes as these can be interpreted as specific markers of that clone. We used MAST^61^ with log_2_-normalized counts (pseudocount=1) as input and cellular detection rate as covariate. P-values were computed by likelihood ratio test and then adjusted for multiple testing using the Benjamini-Hochberg correction. We used MAST coefficients (log_2_ fold change) as input to the prerank function of GSEApy^62^ with MSigDB Hallmark gene sets^60^.

#### IFN-γ response gene score

To compute the IFN-γ response gene score (**Fig. 5f,h**; **Supplementary Fig. 10h**), we took the mean of z-scored log-normalized expression across genes in this Hallmark set. All genes in the set were included if they had been retained during highly variable gene selection for the relevant integrated MrVI model.

#### Classifying cycling cells

Cell cycle phase (**Supplementary Fig. 11a,b**) was assigned to each cell using z-scored log-normalized expression as input to the built-in Scanpy function (score_genes_cell_cycle) with established cell cycle genes^76^. The fraction of cycling cells (S or G2M phase) was reported for each cell type, sample, and genotype combination with at least 15-25 cells.

#### Phylogeny reconstruction

Clonal phylogeny was inferred using PICASSO, a probabilistic tool for reconstructing phylogenetic trees from CNV data^44^ **(Fig. 6d**; **Supplementary Fig. 16c,h**). To use PICASSO with mutation data instead of CNVs, we encoded mutations with integer states. First, we filtered variants based on genotype confidence. For each variant, if a sufficient number of cells (1,000-3,000) could be genotyped with high confidence (*P*(*G*) > 0. 9), then these cells were retained, and their genotypes were encoded as 0 (wt/wt), 1 (mut/wt), or 2 (mut/mut). If, instead, cells could be genotyped with only moderate confidence (*P*(*G*) > 0. 6), then these cells were retained, and their genotypes were encoded as 0 (wt/wt) or 2 (mut/wt or mut/mut). Variants that met neither of these criteria were excluded. Cells were then retained if genotyped for all included variants. In some cases, variants were omitted to ensure that enough fully genotyped cells could be retained. The variants included for PICASSO are shown in **Fig. 6a** and **Supplementary Fig. 16b,f** (except *IDH1* c.394C>T).

We ran PICASSO for each patient on the filtered genotype matrix with maximum tree depth of 6 and termination by probability. Minimum clone size was set by patient (4-40 cells). We manually annotated *TP53* LOH (**Fig. 6d**) and *JAK2* LOH (**Supplementary Fig. 15c**) based on mutation frequencies (**Supplementary Fig. 14f**). We also manually appended *JAK2* and *IDH1* mutations to the PICASSO output for **Supplementary Fig. 16g** based on significant mutation co-occurrence (Fisher’s exact test). We assigned cell clusters to clones based on genotype frequencies in each PhenoGraph cluster, excluding those with ambiguous assignments.

#### Copy number inference for transforming MPN patient

We computed per-gene CNVs for clones A, C, and D (**Fig. 6d**) using infercnvpy with a sliding window of 100 genes, step size of 10, and clone A as the reference population. Log-normalized counts for all genes (pre-filtering) were used as input. Mean per-gene CNV values were calculated for each clone and plotted against the chromosomal gene start position (**Supplementary Fig. 15**).

#### Determination of possible lineage trees for GBM patient

To determine the number of possible lineage trees compatible with the bulk mutations observed in the GBM patient shown in **Fig. 3**, we first constrained the mutation set to the 4 variants shown in **Fig. 3f,g**. We define a tree as distinct if it yields a unique set of terminal states, or sub-clones, as visualized for the set of trees in **Supplementary Fig. 5j**. We assume that all intermediate subclones are retained and that all lineages branch from a common wildtype ancestor. Further, we assume an infinite sites model.

There are five possible structures for mutation acquisition in this setup:

1. 4 lineages (*n* = 1): Each mutation arises in an independent lineage, yielding four subclones each with one mutation.
2. 3 lineages (*n* = 12): One lineage acquires two mutations, and two lineages acquire one mutation. There are 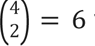 ways to select the pair of mutations in the same lineage, and there are 2 internal orderings for each pair (e.g. mutations A then B yields sub-clones A and AB; B then A yields sub-clones B and AB).
3. 2 lineages with 3 mutations in 1 lineage (*n* = 24): One lineage acquires three mutations, and one lineage acquires one mutation. There are 4 ways to assign the single mutation, and there are 3! possible orderings of 3 mutations in one lineage. Thus, 4 * 3! = 24 possible sets.
4. 2 lineages with 2 mutations in each (*n* = 12): There are 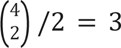 ways to split the mutations into two unordered pairs, and there are 2 * 2 possible orderings per set of pairs. Thus, 3 * 4 = 12 possible sets.
5. 1 lineage with all 4 mutations (*n* = 24): All mutations occur sequentially in a single lineage, so the number of possible trees equals the number of possible orders, or 4!.

The total number of distinct lineage trees, or sub-clone sets, is 1 + 12 + 24 + 12 + 24 = 73.

## CODE AVAILABILITY

Custom code and intermediate data files to reproduce all analyses supporting this manuscript are available at https://github.com/clareaulab/gift_figures_reproducibility. Supporting code for gapped probe design is available at https://github.com/clareaulab/probeset_design_pipeline. Software for parsing Gift sequencing data via giftwrap is available at https://github.com/clareaulab/giftwrap. A full step-by-step protocol for GIFT is available on <u>protocols.io</u>

## DATA AVAILABILITY

Sequencing data associated with this work is available at GEO accession **GSE319999** with reviewer access code **yhepqggyhxyfvcz**.

## SUPPLEMENTARY FIGURES

**Supplementary Figure 1.**
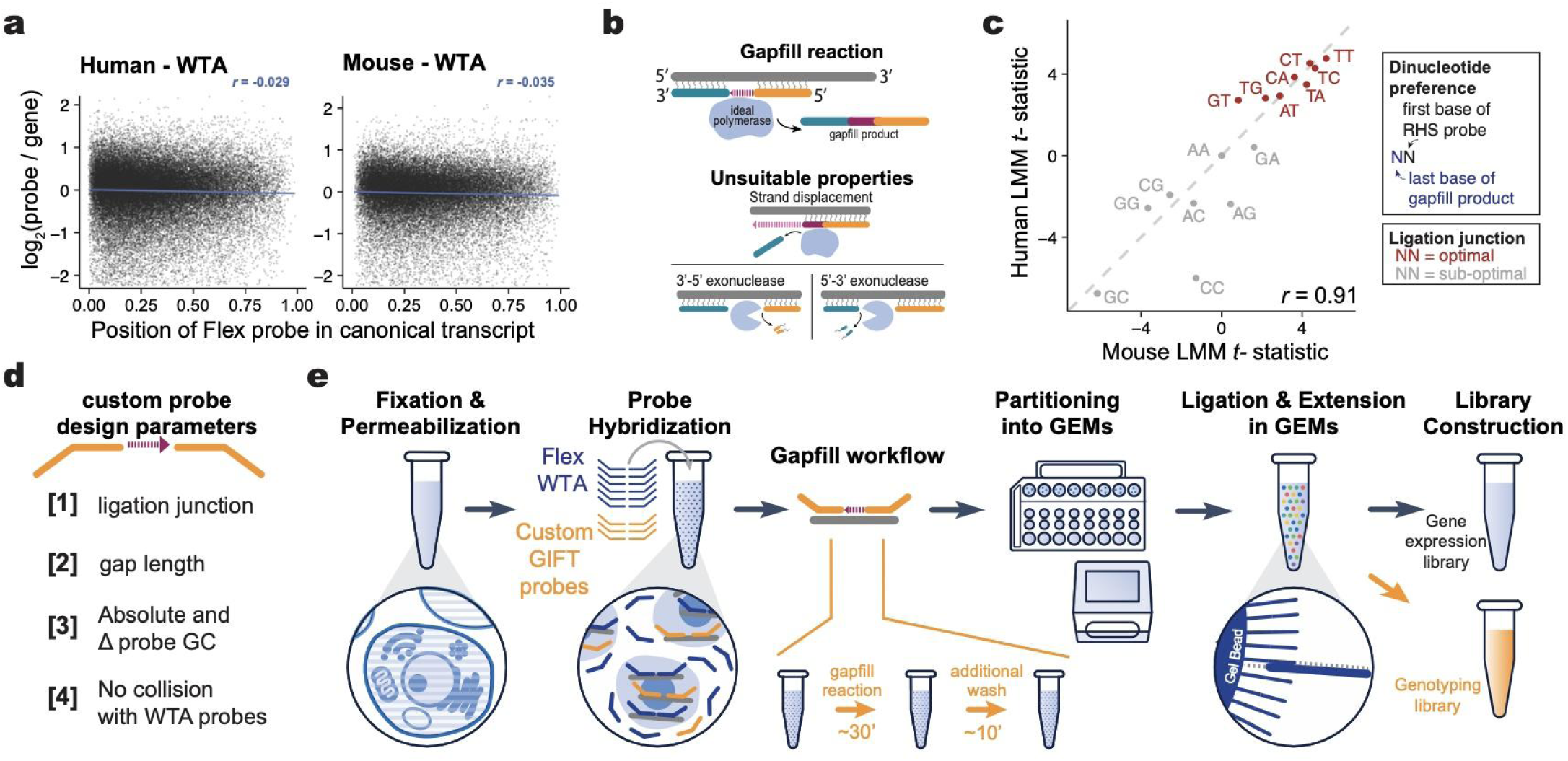
Development of GIFT. **a,** Normalized probe count vs. transcript position for Flex WTA panels showing that probe-based detection is possible throughout the transcript (n=53,330 human; 54,263 mouse probe pairs). **b**, A suitable gapfill polymerase is strand displacement deficient and lacks exonuclease activity. **c**, Summary of linear mixed model (LMM) associations for human and mouse probe sets to define optimal ligation junctions based on the Flex WTA probesets for mouse and human. **d**, Parameters used for custom GIFT probe design. **e**, Detailed overview of flex plus gapfill (GIFT) workflow. Separate PCR handles are used to separately amplify gene expression (WTA) and genotyping (GIFT) libraries.

**Supplementary Figure 2.**
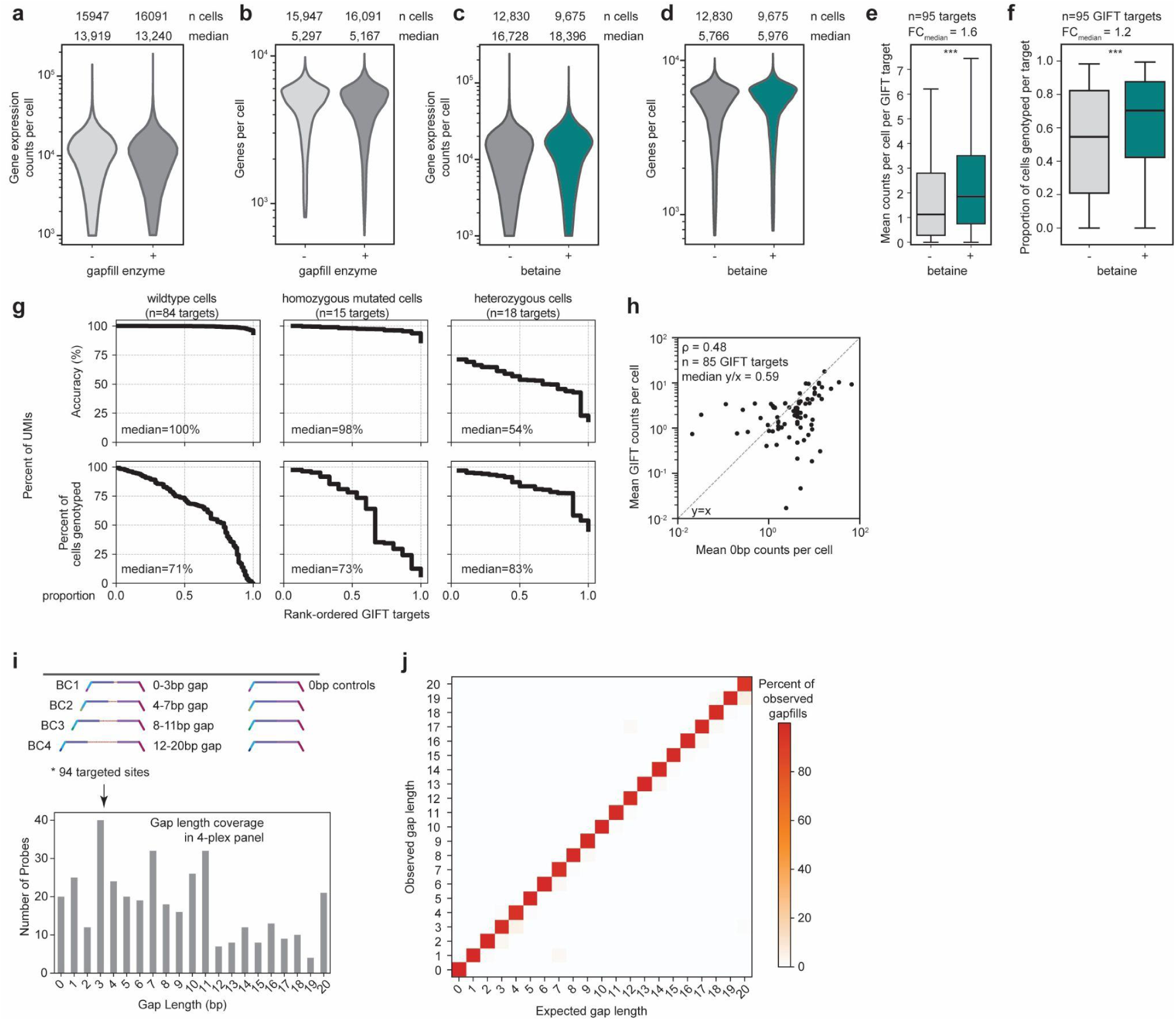
GIFT metrics in cell lines. **a-b**, The number of counts (**a**) or genes (**b**) captured per cell for Flex gene expression assay (WTA) is similar with or without the gapfill polymerase. The median number of counts per cell with gapfill enzyme is 95% of the median per cell without (one-sided Mann-Whitney U; p < 0.0005). The median number of genes captured per cell with gapfill enzyme is 97.5% of the median per cell without (one-sided Mann-Whitney U; p < 0.0005). Other GIFT steps were included for both conditions with only polymerase omitted for “- enzyme”. Sequencing saturation was 15% (with enzyme) or 14% (without enzyme). **c-d**, The number of counts (**c**) or genes (**d**) captured per cell for gene expression (WTA) is similar with or without betaine. The median number of counts per cell with betaine is 110% of the median per cell without (p < 0.0005, one-sided Mann-Whitney U test). The median number of genes captured per cell with betaine is 104% of the median per cell without (p < 0.0005, one-sided Mann-Whitney U test). Sequencing saturation was 18% (with betaine) or 16% (without betaine). **e-f**, Genotyping yields for GIFT carried out with or without betaine, reported as mean counts per cell per target (**e**) and proportion of cells genotyped per target (**f**, ≥1 count per cell). Betaine increases yield (p < 0.0005, one-sided Wilcoxon test). The same GIFT targets were used for both experiments. **g**, Cell genotyping metrics across all targets in the cell line mixing experiment shown in Fig. 1. *Top*: Genotyping accuracy for genotyped cells (≥1 GIFT count). *Bottom*: Proportion of cells for which GIFT assigns a genotype. **h**, GIFT counts per cell versus gene-matched 0-bp control probes (ρ, Spearman correlation; y/x measures gapfill efficiency). Control probes are targeted elsewhere on the same transcript but have no gap between probes. **i**, Design of gap length experiment. *Top*: Probe design strategy for a representative targeted site. The gap lengths for each probe barcode varied by target; for example, some targets had the longest gap on BC1 or BC2 or BC3 rather than BC4 as depicted for the target shown. *Bottom*: Number of probes for each gap length. Full panel is in **Extended Data Table 1b**. **j**, Observed vs. expected gap lengths for experiment described in **i**.

**Supplementary Figure 3:**
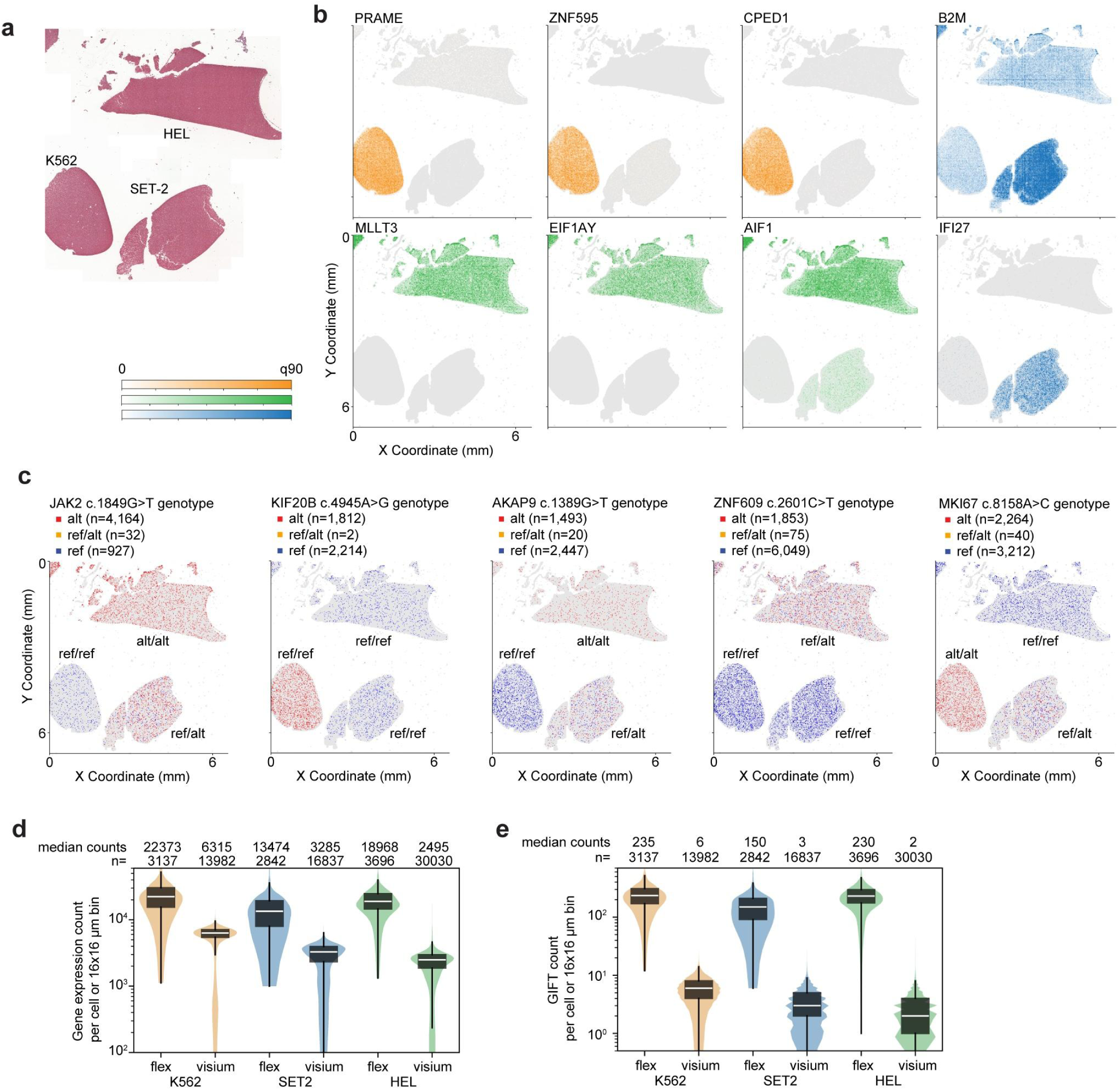
GIFT in Visium HD. **a**, H&E staining for spatial GIFT showing locations of cells in the capture area. **b**, Cell type-specific marker gene expression confirming cell identity for the patterning of the cell lines. q90, 90^th^ percentile of normalized counts per bin. **c,** Spatial genotyping showing the captured alleles in each 16 x 16 μm bin. Expected genotypes by cell line are annotated next to the spatial region. **d-e**, Comparison of gene expression (**d**) and GIFT genotyping (**e**) yields per cell or 16 x 16 μm bin from Flex or Visium HD, respectively, stratified by cell line. The same GIFT targets were included for both experiments (n=84 shared targets).

**Supplementary Figure 4:**
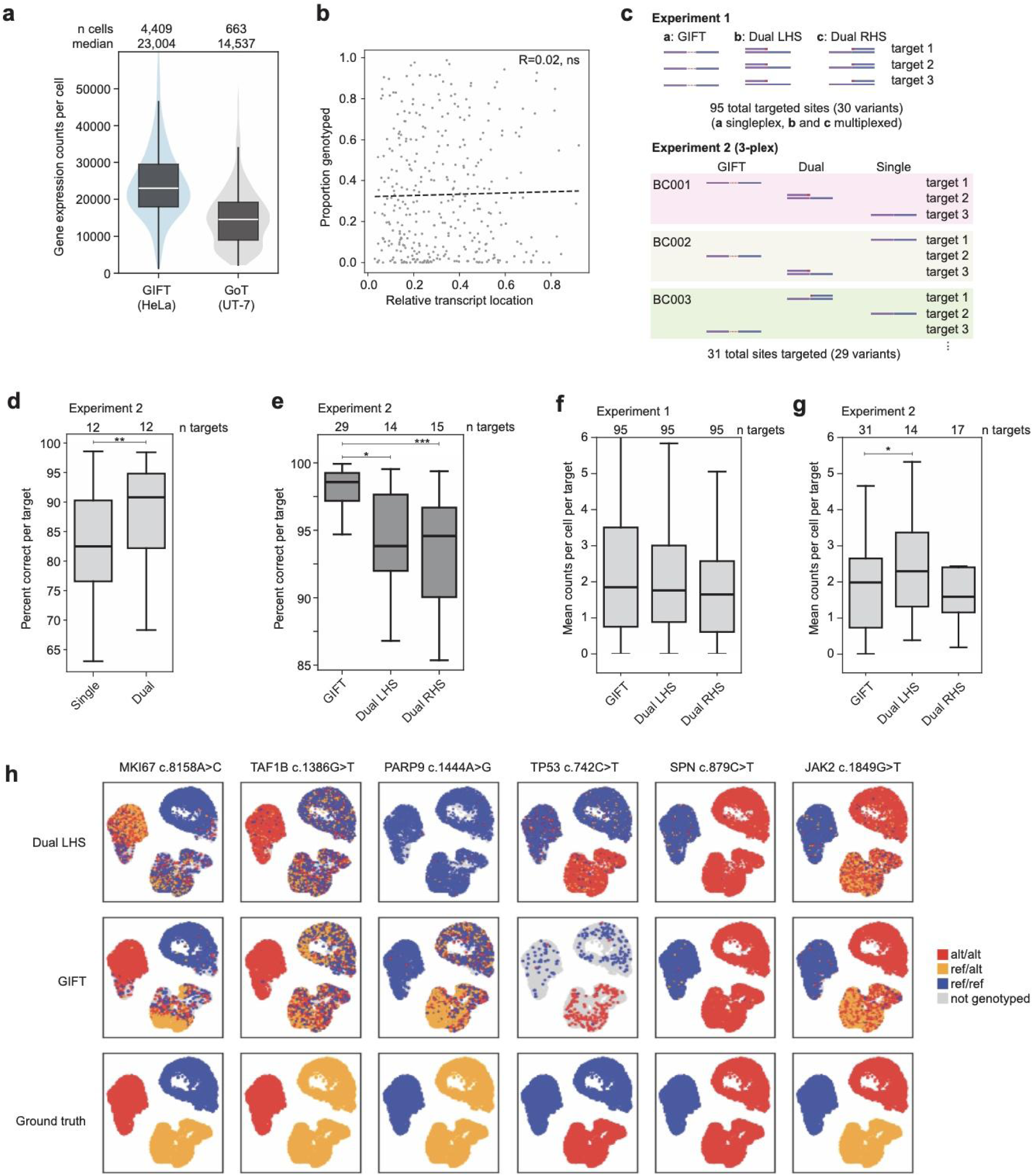
GIFT benchmarking in cell lines. **a-b**, Comparisons between GIFT (this work) and GoT^10^. **a**, Gene expression counts per cell for GIFT vs GoT for 17,225 genes captured in both experiments. **b**, GIFT genotyping performance is not dependent on transcript location (R, Pearson correlation; p-value=0.7, n=344 GIFT targets). **c-h**, Benchmarking of GIFT (gapfill) against dual probe and single probe genotyping. **c**, Overview of two experiments. Genotyping accuracy for experiment 1 is shown in Fig. 2f**,g**. **d-e**, Percent of genotyping counts (UMIs) that are correct for each target by method (Dual vs. Single probe genotyping in **d** and GIFT vs. Dual in **e**). Targets are shared between GIFT and Dual probes but are mutually exclusive for Dual LHS and Dual RHS (*** p < 0.0005, ** p < 0.005, * p < 0.05, one-sided Wilcoxon test). **f-g**, Mean counts per cell per target by genotyping strategy for each experiment (* p < 0.05, one-sided Wilcoxon test). **h**, UMAP embeddings of representative variants, colored by Dual LHS, GIFT, or ground truth (bulk) genotypes (n = 9,675 GIFT-assayed cells, n = 16,097 Dual probe-assayed cells).

**Supplementary Figure 5:**
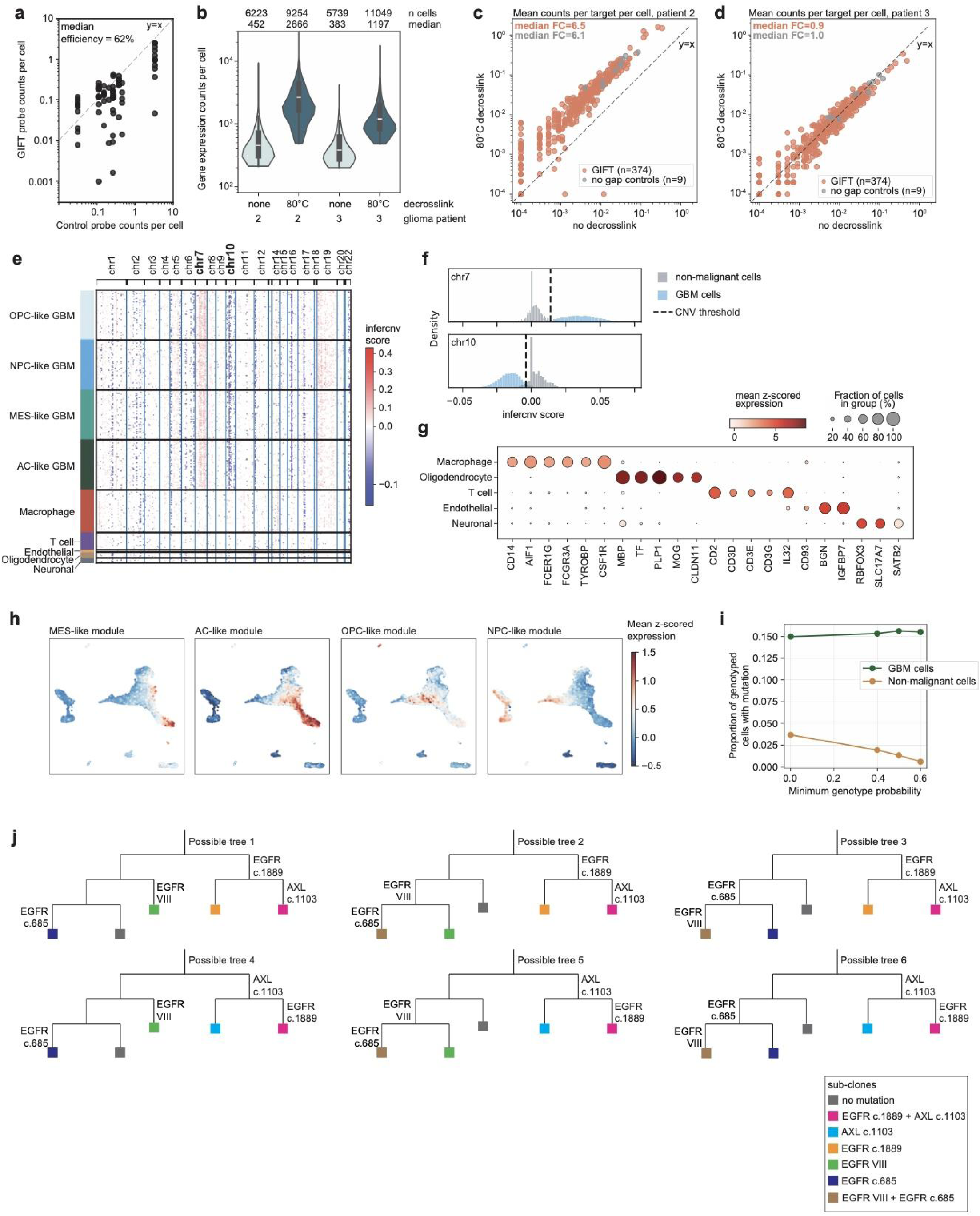
GIFT genotyping in archival FFPE tissue. **a**, GIFT counts per cell versus gene-matched 0-bp control probes for FFPE GBM tissue (80° dcl, n=72 GIFT targets with matched controls), as shown for cell lines in **Supplementary** Fig. 2h. This is for the same patient as in Fig. 3, hereafter called patient 1. Gapfill efficiency is the ratio of GIFT to control counts (y/x). Control probes are targeted elsewhere on the same transcript but have no gap between probes. **b-d**, Effect of decrosslinking on gene expression (WTA, **b**) and GIFT genotyping (**c**-**d**) yields in 2 additional glioma patients. Decrosslinking improves gene expression yield per cell for all patients. Decrosslinking improves GIFT yield per cell for patient 2. For patient 3, decrosslinking does not improve GIFT yield per cell but also does not improve yields for the 0-bp control probes, despite improvements for WTA. **e**, Inferred CNVs for GBM patient 1. GBM cells show gain of chromosome 7 and loss of chromosome 10. NPC, neural progenitor cell; AC, astrocyte; MES, mesenchymal; OPC, oligodendrocyte progenitor cell. **f**, Distributions of infercnv scores for chromosome 7 (*top*) and chromosome 10 (*bottom*) for GBM and non-malignant cells. Thresholds were added by visual inspection and used to classify cells in Fig. 3g. **g**, Z-scored, log-normalized expression of marker genes in non-malignant cell types. **h**, Expression of GBM subtype modules^18^. NPC, neural progenitor cell; AC, astrocyte; MES, mesenchymal; OPC, oligodendrocyte progenitor cell. **i**, Proportion of genotyped cells with at least one mutation based on genotype calls at different probability thresholds. As the minimum genotype probability increases, the false positive rate of mutation in non-malignant cells decreases. The genotype probability is defined as the probability that a cell is homozygous mutated, heterozygous, or wildtype, but homozygous and heterozygous mutation calls are included when calculating the proportion mutated. **j**, Expansion of possible phylogenies encompassed by the minimal tree shown in Fig. 3h. Our assumptions for defining possible lineage trees are detailed in methods.

**Supplementary Figure 6:**
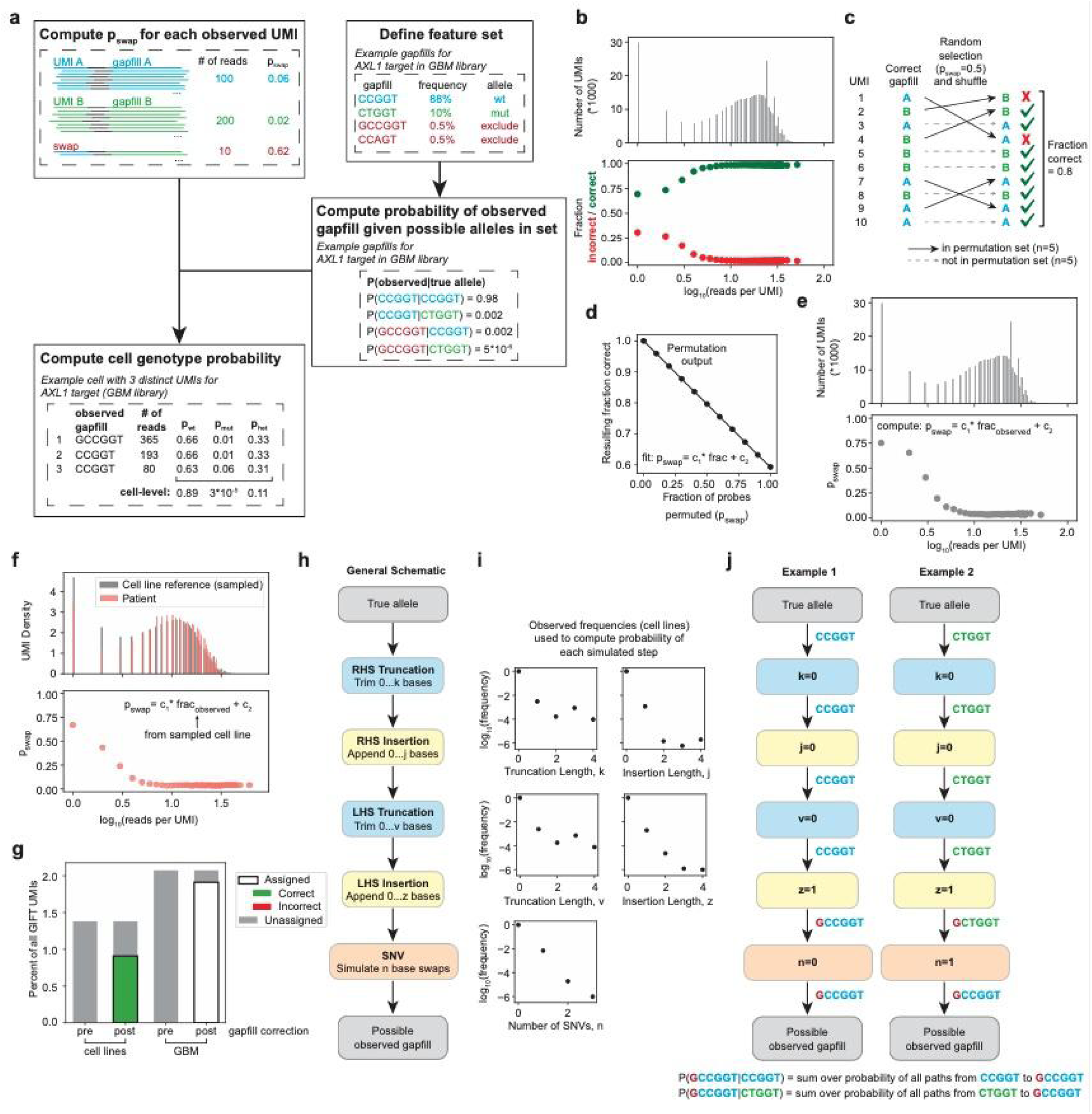
Probabilistic model for assigning genotypes from GIFT. **a**, Our probabilistic strategy for assigning genotypes from GIFT counts (**methods**). In the first step, we use the number of reads per UMI to compute the probability (*p_swap_*) that a UMI is an artifact of PCR-mediated template switching. An example of a swapped product with few reads is shown. In the second step, we define a constrained feature set that excludes observed gapfills without significant signal above noise. In the third step, we compute for each observed gapfill the probability of observation given possible true alleles in the set. For gapfills that are not included in the set, such as the example shown in red, it is still much more likely to come from one possible allele than the other. In the fourth step, we integrate information across UMIs for the same target in each cell to compute probabilities that a cell is wildtype, homozygous mutated, or heterozygous. An example calculation for a cell with 3 observed *AXL1* counts is shown. **b-f**, Details of *p_swap_* calculation. **b**, *Top*: Distribution of reads per UMI for a representative cell line sample. *Bottom*: Probability that a UMI is correct (matches ground truth genotype) or incorrect as a function of reads per UMI. **c**, Permutation approach used to infer *p_swap_* from the observed fraction correct. We took all UMIs in the dataset and shuffled the genotypes of a random selection of UMIs. The proportion of UMIs selected for shuffling is a proxy for *p_swap_*. This example shows a set of 10 UMIs permuted at *p_swap_* = 0.5. **d**, Linear regression relating *p_swap_* to the resulting fraction correct from the synthetic permutations. This equation can then be used to compute *p_swap_* from the observed fraction correct in a real dataset. **e**, *Top*: Distribution of reads per UMI in the cell line dataset as in **b**. *Bottom*: Inferred *p_swap_* computed from the fraction correct values in **b** and the regression fit in **d**. **f**, Inference of *p_swap_* for patient data using the values computed for the cell line data. The cell line reference is sampled to closely match the distribution of reads per UMI for the patient dataset, which accounts for differences in sequencing depth. Then, the cell line fraction correct at each sampled read count is used to compute *p_swap_* for the patient UMIs. **g-j**, Details of edit distance calculations used to compute the probability of an observed gapfill given possible true alleles. **g**, Breakdown of GIFT UMIs requiring gapfill correction, which is detailed in subsequent panels. Before gapfill correction, unassigned UMIs are any gapfills that do not exactly match an allele in the feature set. After gapfill correction, assigned UMIs are those for which we can assign an allele with >90% confidence. In cell lines, we confirm that accuracy in this set is high (99.8% correct). **h-i**, Simulated steps to model the possible alterations in a gapfill sequence that could occur from the true allele template to the observed sequence (**h**). These changes could occur during the gapfill polymerase step, PCR, or sequencing, and we determine their rates empirically based on observed gapfill sequences in cell line datasets (**i**). We compute the probability of each observed sequence using the product of the inferred probabilities at each step. **j**, Examples of edit distance calculation for an observed gapfill (GCCGGT) and two possible true alleles in the feature set. Example 1 (CCGGT) requires fewer changes to the original sequence, which is why this is much more likely to be the true allele, as shown by the calculation of P(observed|true allele) in **a**.

**Supplementary Figure 7:**
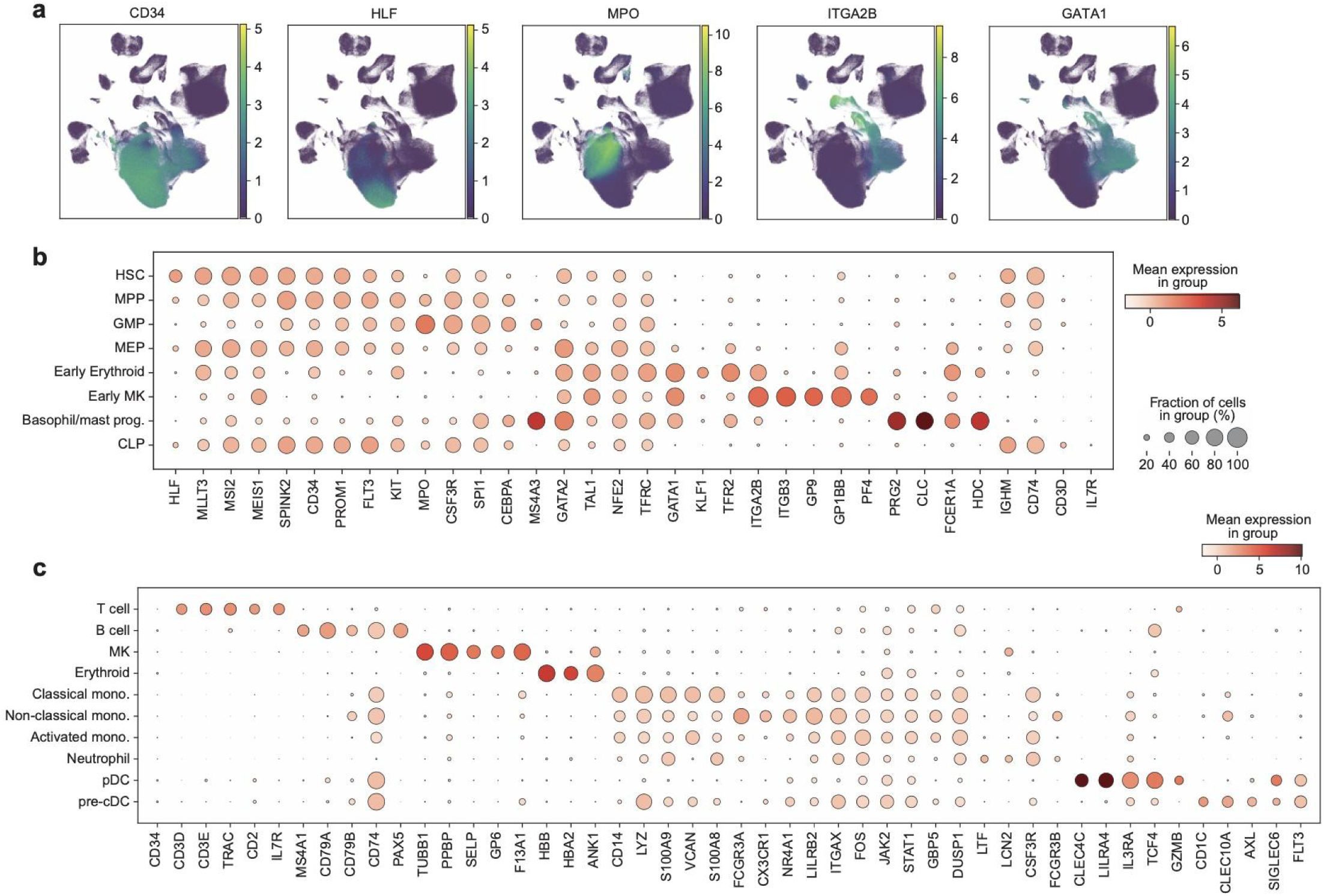
Markers for cell typing in MPNs. **a**, UMAPs of patient-integrated scVI^59^ latent space colored by log-normalized expression of selected hematopoiesis markers^28^. **b**, Z-scored log-normalized expression of selected markers of HSPC cell types. HSC, hematopoietic stem cell; MPP, multipotent progenitor; GMP, granulocyte-monocyte progenitor; MEP, megakaryocyte–erythroid progenitor; MK, megakaryocyte; CLP, common lymphoid progenitor. **c**, Z-scored log-normalized expression of selected markers of mature cell types. MK, megakaryocyte; pDC, plasmacytoid dendritic cell; cDC, conventional dendritic cell.

**Supplementary Figure 8:**
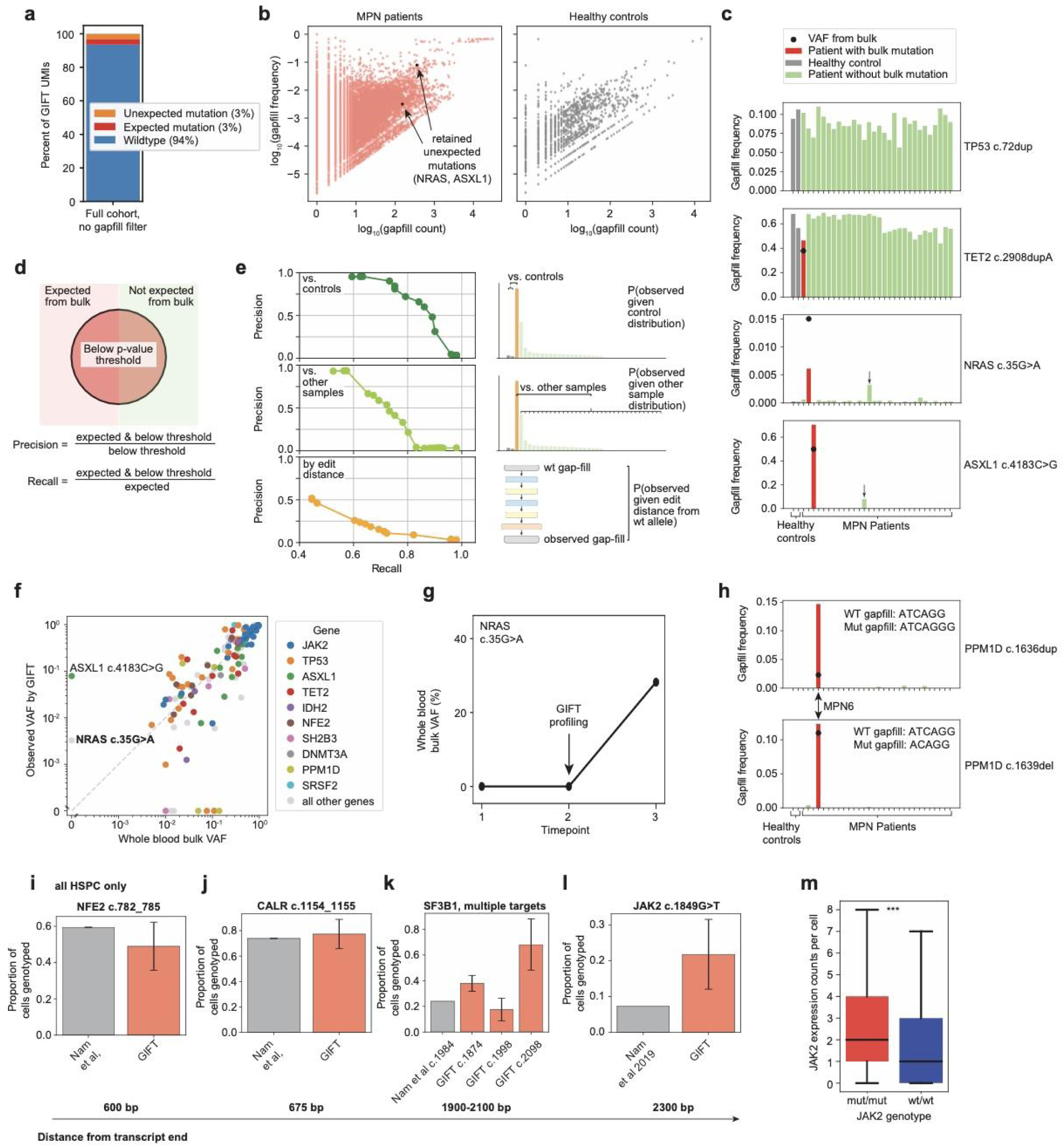
Variant discovery and genotyping flexibility with GIFT in MPN cohort. **a**, Overview of observed gapfill sequences in the MPN cohort. Expected mutations are UMIs with gapfills exactly matching expected mutated alleles. Unexpected mutations are UMIs with gapfills that match neither wildtype nor expected mutation alleles. **b**, Frequency and count of gapfills with unexpected mutations in the MPN cohort. Gapfill count is the number of UMIs of that gapfill per patient, and gapfill frequency is the count normalized by the total observed UMIs for that targeted site (any gapfill sequence) for that patient. *Left*: all unexpected gapfills (n=38,344 across all patients). *Right*: unexpected gapfills observed in healthy controls (n=2,988) to show scale of background noise. Sequencing saturation of GIFT for all patients is >95%. **c,** Representative observed gapfills and their frequencies across samples. The top two variants (*TP53*, *TET2*) have a high background in healthy controls. The bottom two variants (*NRAS*, *ASXL1*) were expected in two patients but also were significantly enriched in two other patients (arrows). **d-e**, Defining an empirical feature set using observed gapfills in the MPN cohort. **d**, Precision and recall definitions for testing how well each model distinguishes true variants (expected mutations) from noise (vast majority of unexpected gapfills). **e**, Precision-recall curves for three different models, which are detailed in methods, that can be used to assign the probability of observing each gapfill. As shown by the schematics on the right, the control model compares the observed gapfill frequency to that in healthy controls, the other sample model compares the observed gapfill frequency to that in other samples, and the edit distance model compares the observed gapfill frequency to the expected frequency if the true allele is wildtype as calculated in **Supplementary** Fig. 6h**-j**. **f**, VAF observed by GIFT vs. bulk whole blood, as in Fig. 4f but on a log scale. Labeled variants were not expected from bulk sequencing but were discovered as significant under the models shown in **e** using thresholds described in methods. In contrast, four variants were expected at VAF >= 0.05 but were not observed (2 *TP53*, 2 *PPM1D*). Three of these targets were missed because of low detection (probe counts <15), while one (*TP53* c.733G>A) was confirmed to have a lower VAF (0.01) in CD34+ cells than in whole blood (**Extended Data Table 2**). **g**, Longitudinal whole blood bulk VAF of *NRAS* c.35G>A showing that this mutation was not detected at the time of GIFT profiling but becomes prevalent in this patient at a subsequent timepoint. **h**, Frequencies of *PPM1D* mutations that can be captured by the same probe pair and gap. **i,** Proportion of cells genotyped by GIFT vs GoT (Nam et al)^10^. Variants or regions (*SF3B1*) are shown that were profiled in both studies. For variants near transcript ends (*NFE2*, *CALR*), GIFT capture is similar to GoT, while GIFT capture is better for variants far from transcript ends (*SF3B1*, *JAK2*) (GoT n=1; GIFT n=3, 14, 3, 21, 21, 36 profiled patients). Only HSPCs are included in calculations. **m**, *JAK2* expression in mutated vs wildtype cells. *JAK2* expression in *JAK2* V617F^mut/mut^ cells is significantly higher than *JAK2* V617F^wt/wt^ cells (n=138,090 mut cells, 91,513 wt cells, p < 0.0005, one-sided Mann-Whitney U).

**Supplementary Figure 9:**
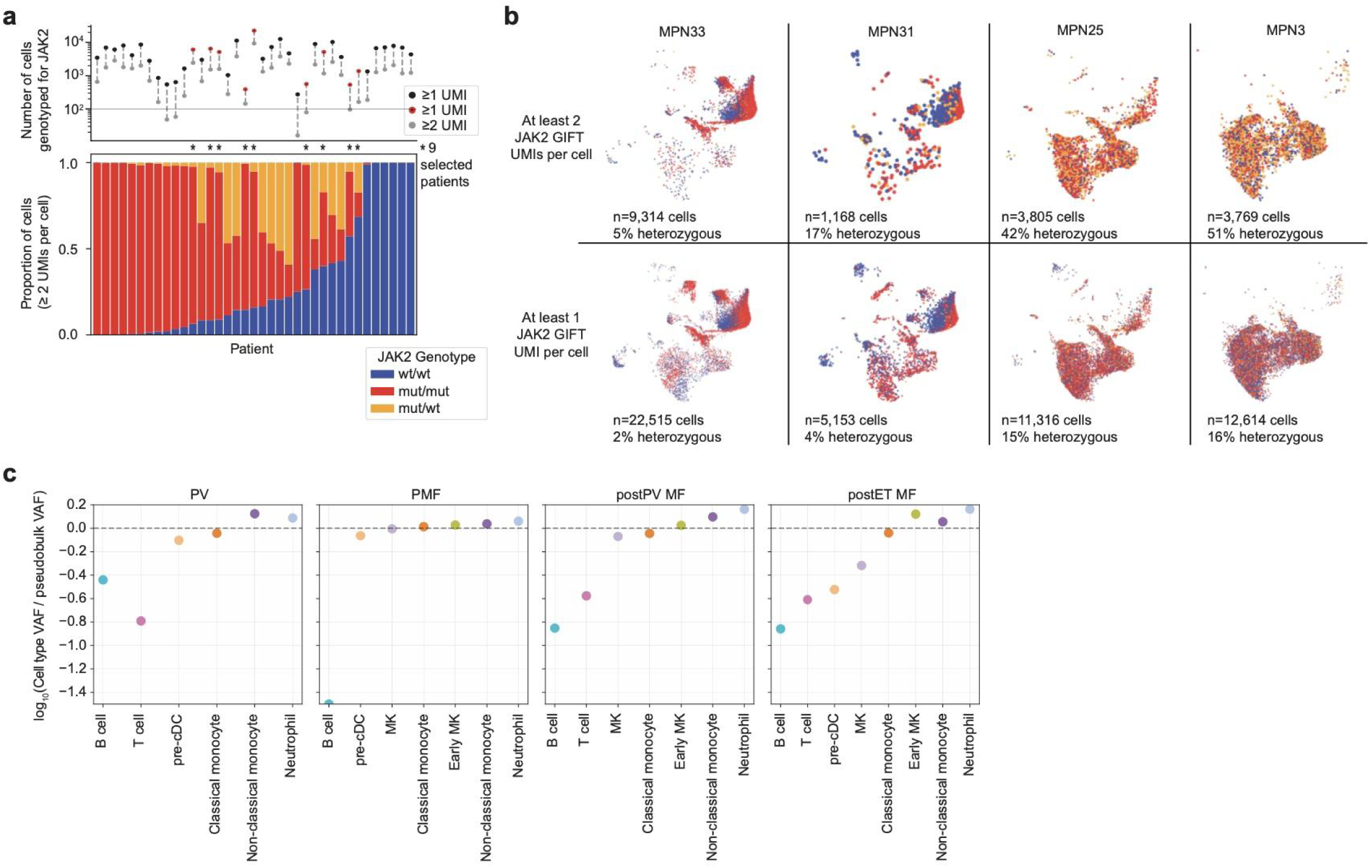
JAK2 genotyping in MPNs. **a**, **a**, *Top*: Number of cells genotyped for *JAK2* V617F when including all cells with ≥ 1 UMI (*black* or *red*) or ≥ 2 UMIs (*grey*) for *JAK2* genotype. Red dots indicate patients selected as primarily homozygous (asterisks on bottom) and included in the MrVI integration used in Fig. 5a**-g**. *Bottom*: Proportion of cells from each patient that are called as homozygous wt (*wt*/*wt*), homozygous mutated (*mut*/*mut*), or heterozygous (*mut*/*wt*) for *JAK2* V617F including only cells with ≥ 2 *JAK2* UMIs. As shown on top, the number of genotyped cells varies widely by patient and is substantially reduced when requiring ≥ 2 UMIs versus ≥ 1 UMI (n cells per patient with ≥ 2 UMIs = 16 - 9,314 cells; n cells per patient with ≥ 1 UMI = 275 - 22,515 cells). **b**, JAK2 genotyping in 4 representative patients. Patients 1 and 2 are primarily wildtype or homozygous mutated so they are included in the patient set in Fig. 5a**-g**. Cells from each patient are shown on the UMAP from the patient-integrated scVI^59^ latent space. **c**, Differential abundance of JAK2 V617F genotype by diagnosis. As in Fig. 5c, the median enrichment of the *JAK2* V617F mutation is shown for each cell type (normalized by pseudobulk VAF across all cell types), but here the values are split by patient diagnosis (n=1-3 patients per datapoint).

**Supplementary Figure 10:**
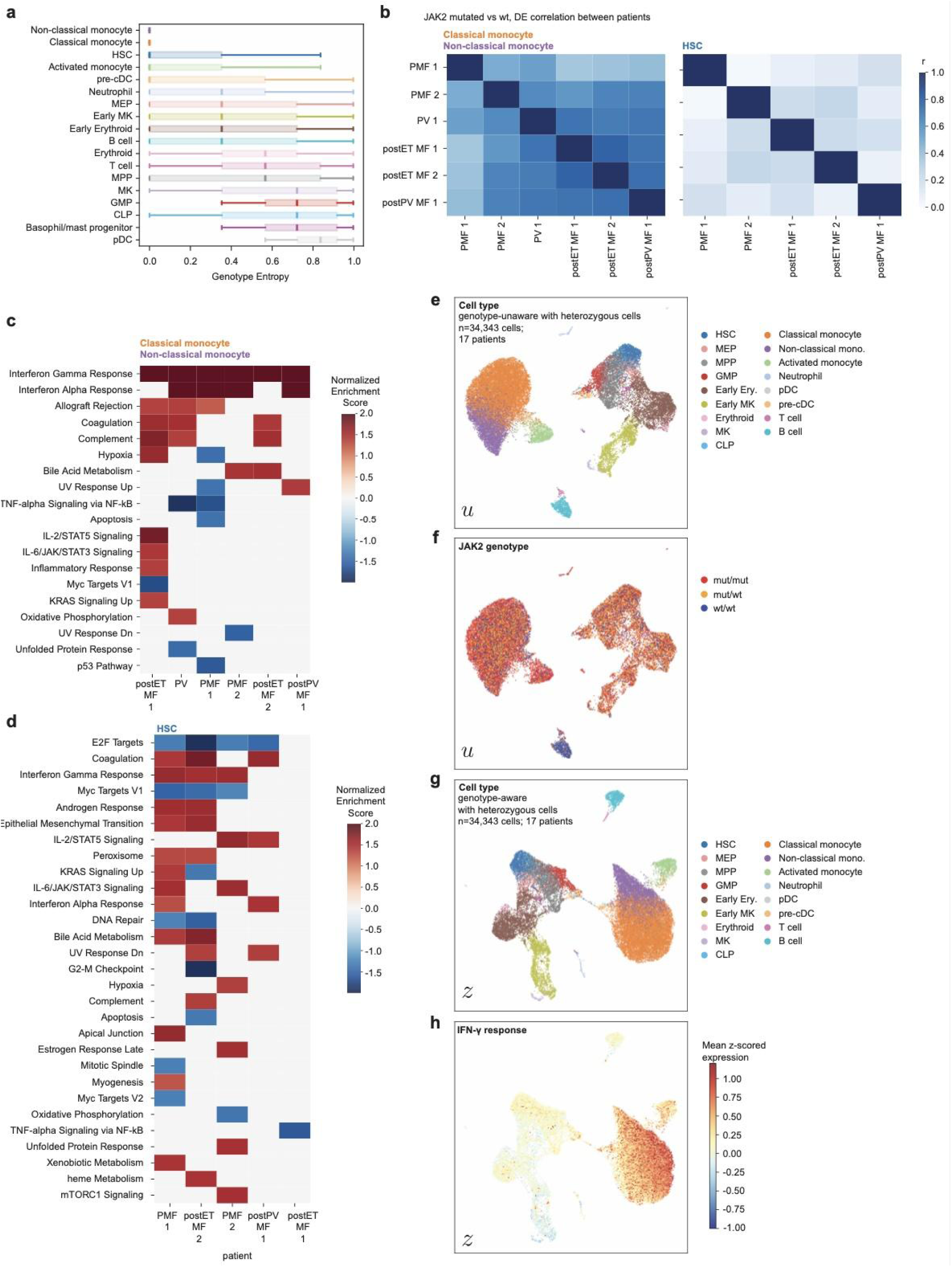
*JAK2*-correlated differential expression across patients. **a,** Genotype entropy in MrVI genotype-aware *z* latent space, stratified by cell type. **b**, Pearson correlation matrices of differential gene expression (log fold-changes, *JAK2*^mut/mut^ vs *JAK2*^wt/wt^) across patients included in Fig. 5g. Each row/column is a patient and is labeled by diagnosis. **c-d**, Differentially expressed pathways by patient for the patients included in Fig. 5g. **e-h**, Additional UMAPs of MrVI latent spaces for the patient set that includes heterozygous *JAK2* mutation (cells in Fig. 5h). **e-f**, Genotype-unaware MrVI latent space (*u*) colored by cell type (**e**) or *JAK2* V617F genotype (**f**). **g-h**, Genotype-aware MrVI latent space (*z*) colored by cell type (**g**) or IFN-γ response gene score (**h**).

**Supplementary Figure 11:**
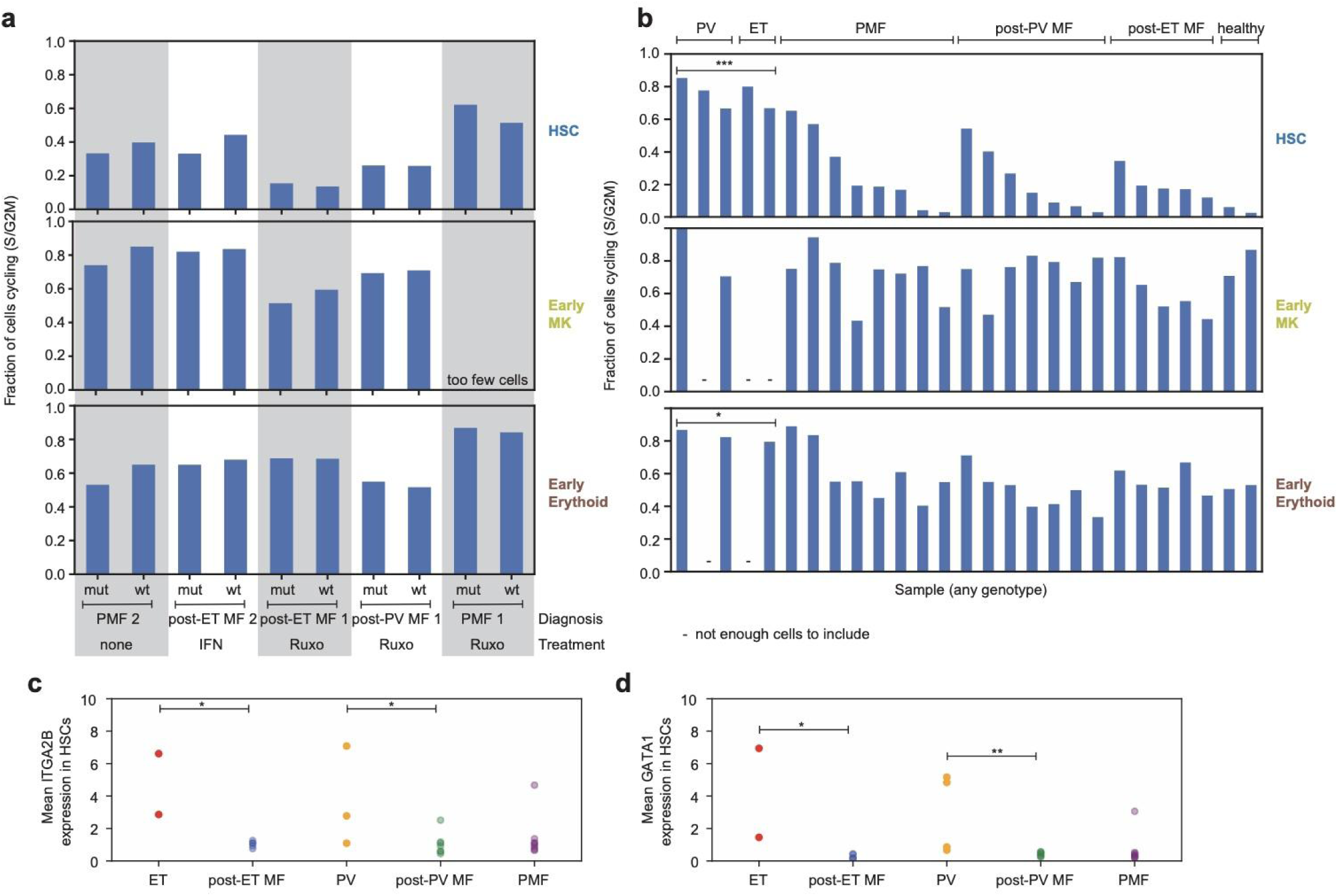
HSPC proliferation and differentiation across MPN types. **a**, Quantification of cell cycling^76^ in patients included in Fig. 5g**-ii**. Little difference is seen in the fraction of cells cycling (categorized as S or G2M phase) for JAK2^mut/mut^ vs JAK2^wt/wt^ cells. **b**, Fraction of cells cycling across all PV (polycythemia vera), ET (essential thrombocythemia), primary myelofibrosis (PMF), post-PV MF (post-polycythemia vera myelofibrosis) and post-ET MF (post-essential thrombocythemia myelofibrosis) patients. Healthy controls are shown for comparison. Patients with PV or ET have higher levels of cycling HSCs and early erythroid cells than MF patients (n=3-5 PV or ET, n=20 MF; one-sided Mann-Whitney U test; * p<0.05, *** p<0.0005). *JAK2* genotype is not considered for this broader patient set because patients may not have enough wildtype or mutated cells, and/or they have too many heterozygous cells, which confounds the analysis. **c-d**, Expression of megakaryocyte (*ITGA2B*) and erythroid (*GATA1*) markers in HSCs suggest greater differentiation bias in ET and PV HSCs than in secondary MF (n=3-8 patients per diagnosis, one-sided Mann-Whitney U test; * p < 0.05, ** p < 0.005).

**Supplementary Figure 12:**
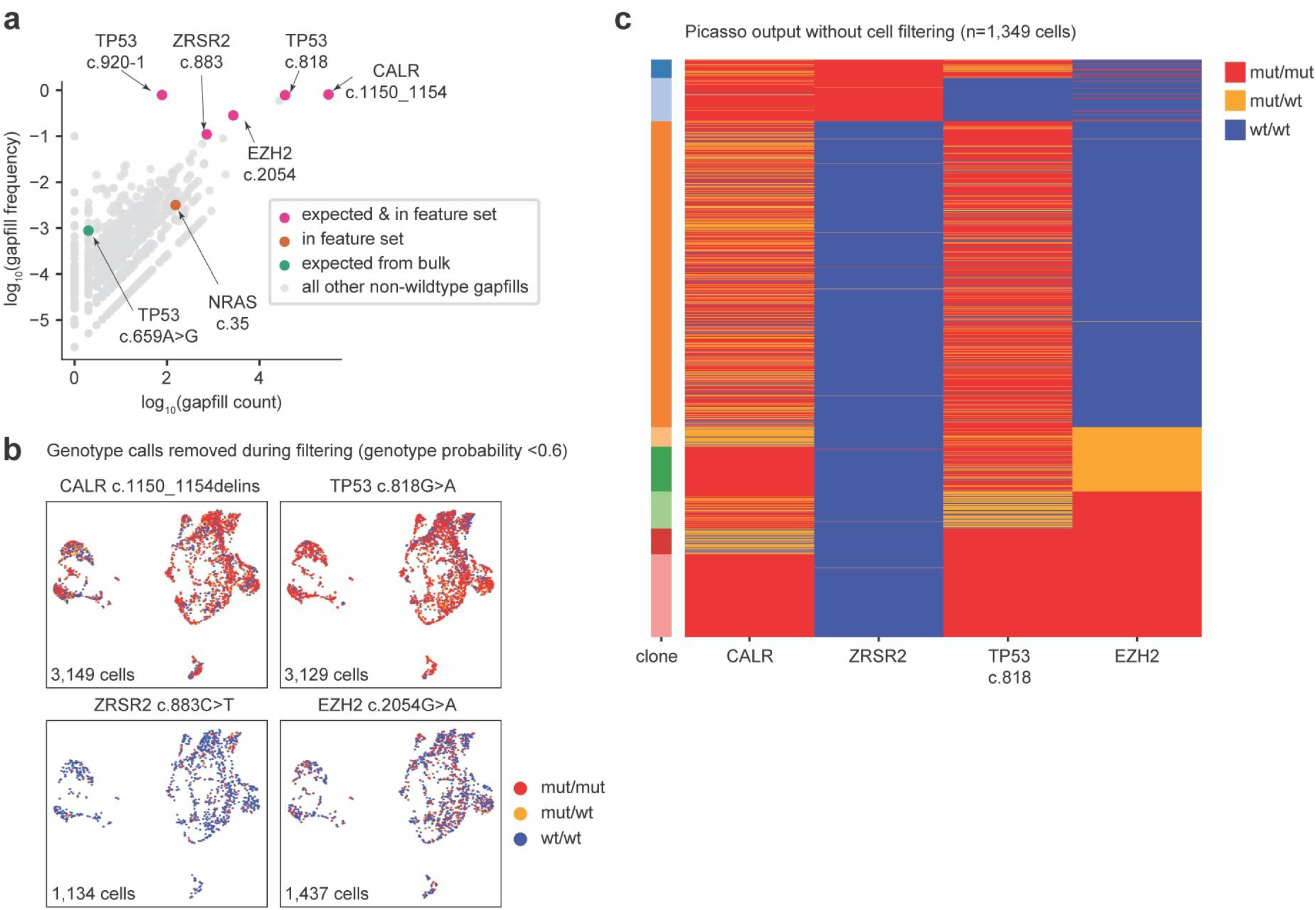
Error model thresholds are necessary for a robust phylogeny. **a**, Frequency and count of gapfills observed in the transforming MPN patient. Frequency is the count normalized by the total observed UMIs for that targeted site (any gapfill sequence). Variants in pink were expected and significantly enriched in GIFT genotyping of this patient. We primarily consider the four of these with the most counts for phylogeny reconstruction. The *NRAS* mutation was discovered by our empirical feature set definition and validated by subsequent bulk sequencing. The *TP53* c.659A>G mutation was expected from whole blood bulk sequencing but not observed by GIFT profiling of the CD34^+^ compartment. Bulk sequencing confirmed a low frequency of this mutation in CD34^+^ cells (**Extended Data Table 2**). The grey dots show observed gapfills that are unlikely to be real mutations and were removed by our feature set definition. **b**, Genotype calls that were removed during filtering [P(genotype) < 0.6). The genotypes in Fig. 6a show the retained set [P(genotype) > 0.6]. Though 0.6 seems like a lenient threshold, the model considers homozygous mutation, heterozygous mutation, and wildtype as possible genotypes, so if P(mut/mut) is ∼0.6, then typically the uncertainty reflects that the cell could be heterozygous [P(mut/wt) >0.3], while the probability that the cell is wildtype is low. **c**, Heatmap of genotypes by clone without probabilistic filtering. This heatmap is generated by PICASSO^44^ as in **Supplementary** Fig. 14a. However, here we include cells genotyped at any confidence level, and we distinguish between heterozygous and homozygous mutation for all variants. Cells with genotypes for all 4 variants are included (n = 1,349 cells). With this unfiltered set of cells, PICASSO finds eight clones, and these are uninterpretable as a meaningful phylogeny.

**Supplementary Figure 13:**
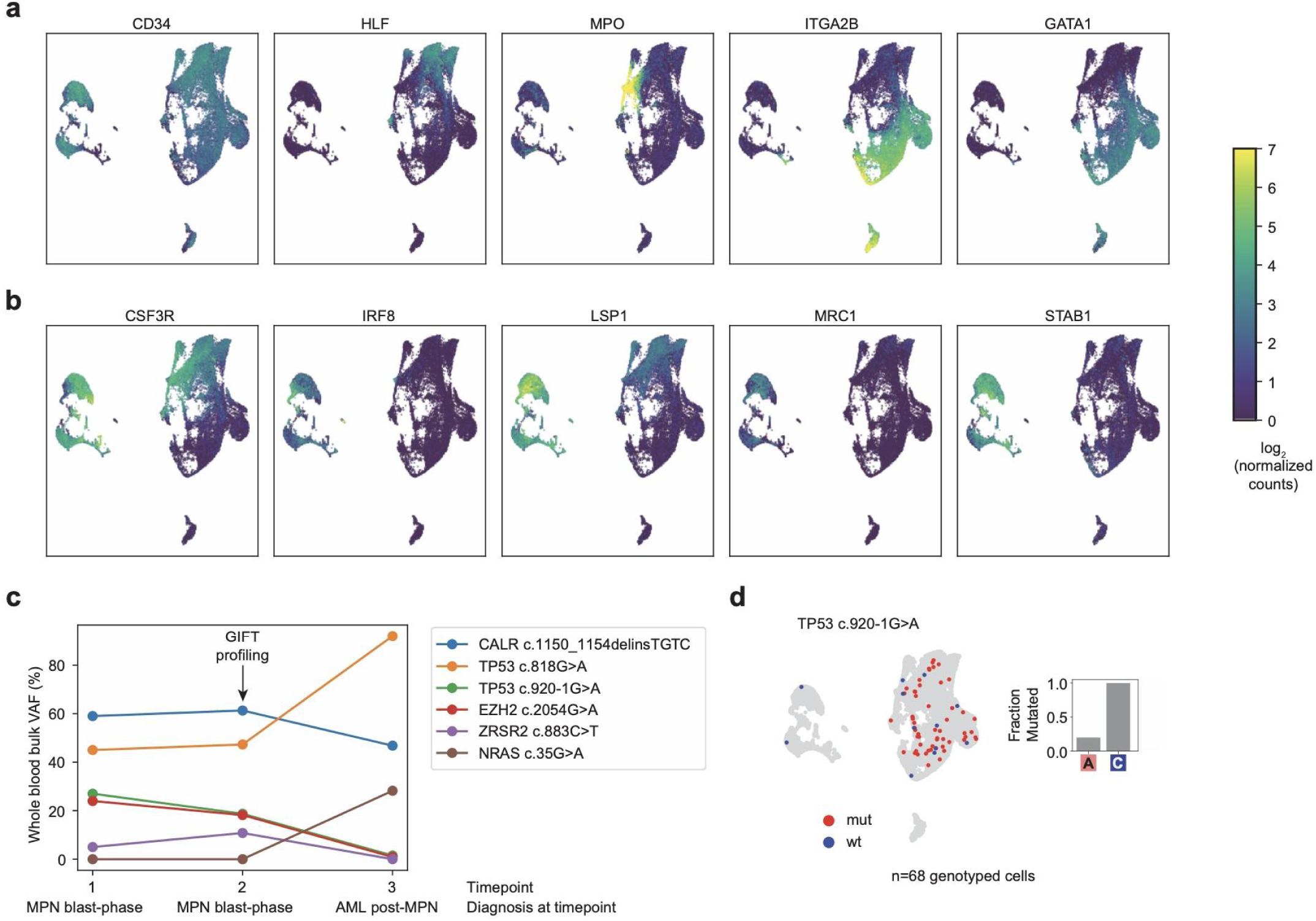
Transformation to AML in an MPN patient. **a**, UMAP embedding of transforming MPN patient as in Fig. 6 but colored by expression of HSPC marker genes^28^. The population labeled as leukemic blast in Fig. 6b expresses *CD34* but not these other canonical HSPC marker genes. **b**, The leukemic blast cells express myeloid markers, matching subsequent diagnosis as acute myeloid leukemia^28,74^. **c**, Whole blood bulk VAFs for this patient at 3 timepoints. The 2nd timepoint was profiled by GIFT. By the 3rd timepoint, the patient’s disease had transformed to post-MPN AML. **d**, Genotyping of splice variant *TP53* c.920-1G>A. The splice variant is captured from pre-mRNA, so detection is low (n=68 genotyped cells), but the mutation is enriched in clone C. This is consistent with disappearance of this mutation at timepoint 3 (as shown in **c**).

**Supplementary Figure 14:**
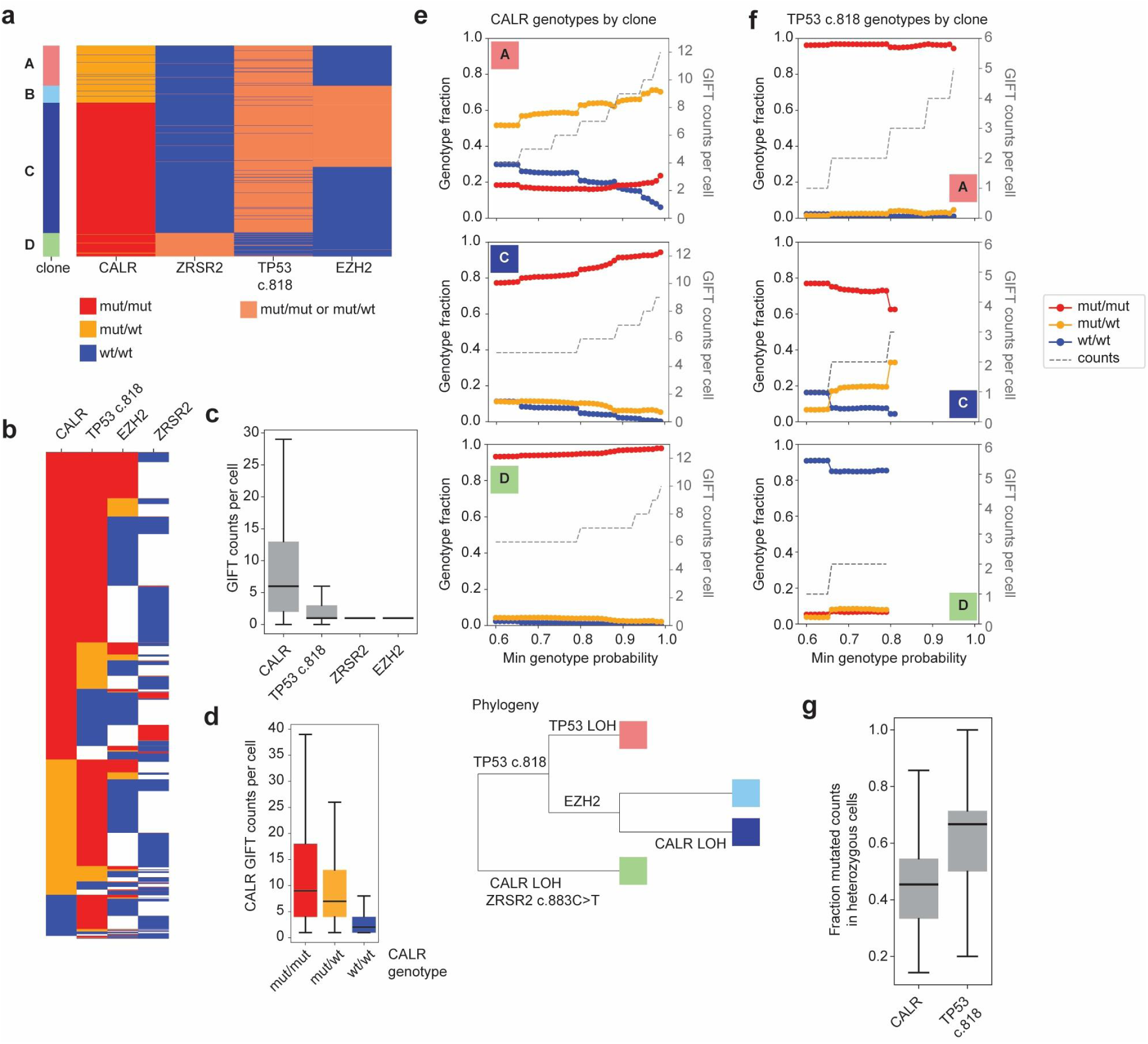
Additional GIFT genotyping details for MPN patient transforming to AML. **a**, Heatmap of genotypes by clone as generated by PICASSO^44^. We had sufficient resolution to distinguish between *CALR* homozygous mutation (mut/mut) and heterozygous mutation (mut/wt) cells. For other variants, we group mut/mut and mut/wt cells together for phylogeny inference by PICASSO. Only cells genotyped for all 4 variants are included (n=730 cells). We manually merged two clones to form clone C because they were artificially separated by *EZH2* wildtype versus mutated calls for cells that are all likely heterozygous. **b**, Heatmap of genotypes sorted by genotype rather than clone and including all cells genotyped for at least 3 variants (n=5,398 cells). Here we distinguish mut/mut and mut/wt calls for all variants to more fully show the underlying data. **c**, GIFT counts per cell by targeted variant (n = 27,180; 16,845; 5,230; 6,894 genotyped cells per variant from left to right). **d**, *CALR* GIFT counts per cell by *CALR* genotype. *CALR* mutation is correlated with higher *CALR* GIFT counts per cell (CALR^mut/mut^ > CALR^mut/wt^ > CALR^wt/wt^; p < 0.0005, one-sided Mann-Whitney U test; n = 3,348-14,726 cells). **e**, *CALR* genotyping by clone at different genotype probability thresholds. Clones B and C are predominantly *CALR*^mut/mut^, especially in high confidence cells. **f**, *TP53* genotyping by clone at different genotype probability thresholds. Cells in clone A are *TP53*^mut/mut^. More cells in clone B are *TP53*^mut/mut^ than *TP53*^mut/het^, but this is biased by higher expression of the mutated *TP53* allele than wildtype allele, as shown in **g**. **g**, Fraction of counts that are mutated for cells that are categorized as heterozygous for *CALR* or *TP53* (most likely genotype by GIFT probabilistic model). Mutated counts are enriched over wildtype counts in *TP53* heterozygous cells (p=1.8*10^-81^; one-sided binomial test, n = 5,337 *TP53*-heterozygous cells; n=62,484 *CALR*-heterozygous cells).

**Supplementary Figure 15:**
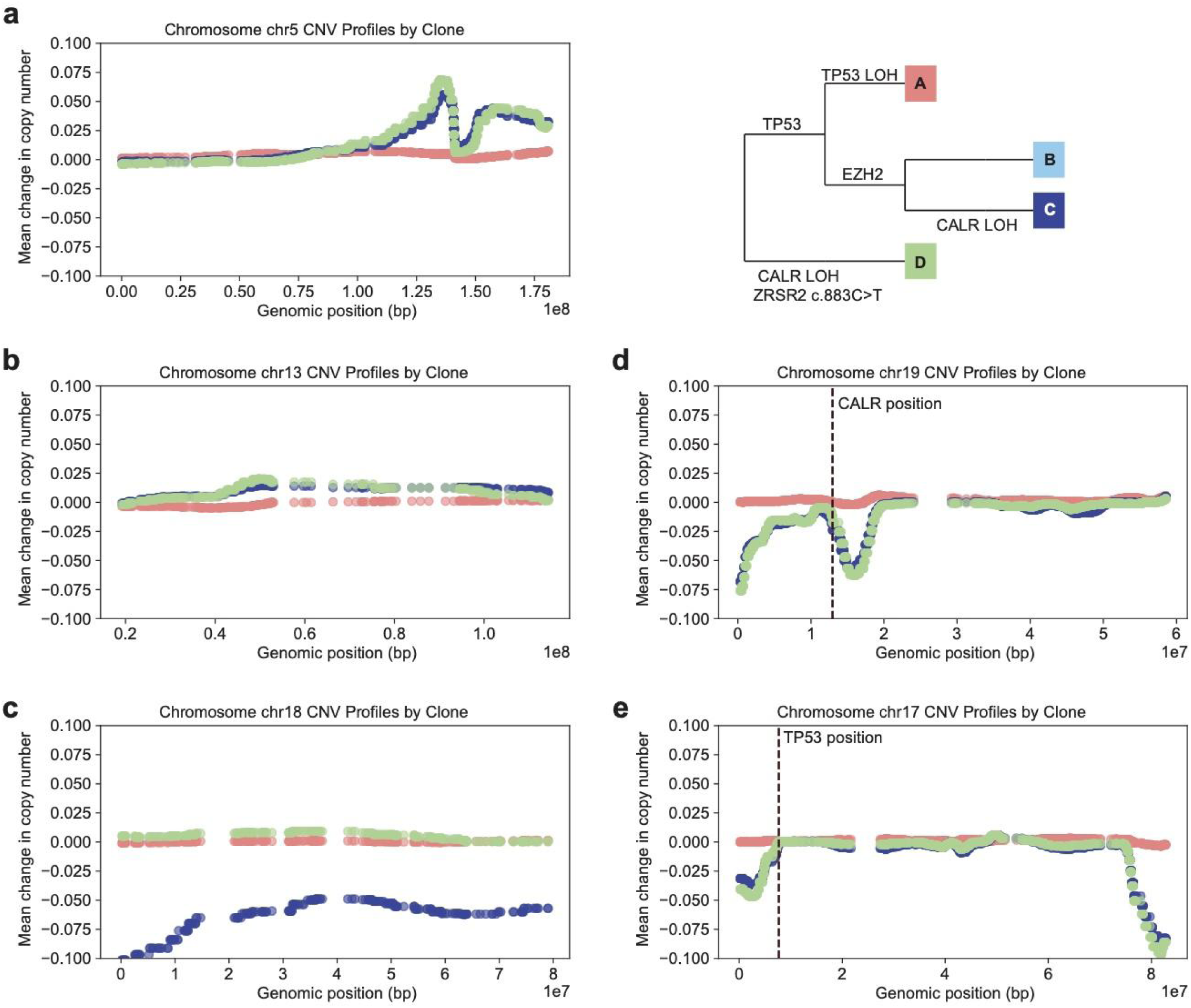
Inferred copy number variants for MPN patient transforming to AML. **a-c**, Inferred CNV profiles from single-cell gene expression. Clone A was used as the copy number reference in all panels. Clone A shows loss of chromosome 5q and chromosome 13q (**a**, **b**), while clone C shows loss of chromosome 18 (**c**). These CNVs were also found independently by clinical karyotyping (**Extended Data Table 2**). **d**, Inferred CNV profile for chromosome 19 showing copy number loss around *CALR* for clones C and D. This is consistent with *CALR* LOH in these clones, as inferred by GIFT genotyping (**Supplementary** Fig. 14e). No CNV was found by clinical karyotyping of chromosome 19 (**Extended Data Table 2**). **e**, Inferred CNV profile for chromosome 17, which contains *TP53*. There is no indication of chromosome loss near TP53 for clone A, despite strong evidence of LOH by genotyping (**Supplementary** Fig. 13c; **Supplementary** Fig. 14f**, Extended Data Table 2**).

**Supplementary Figure 16:**
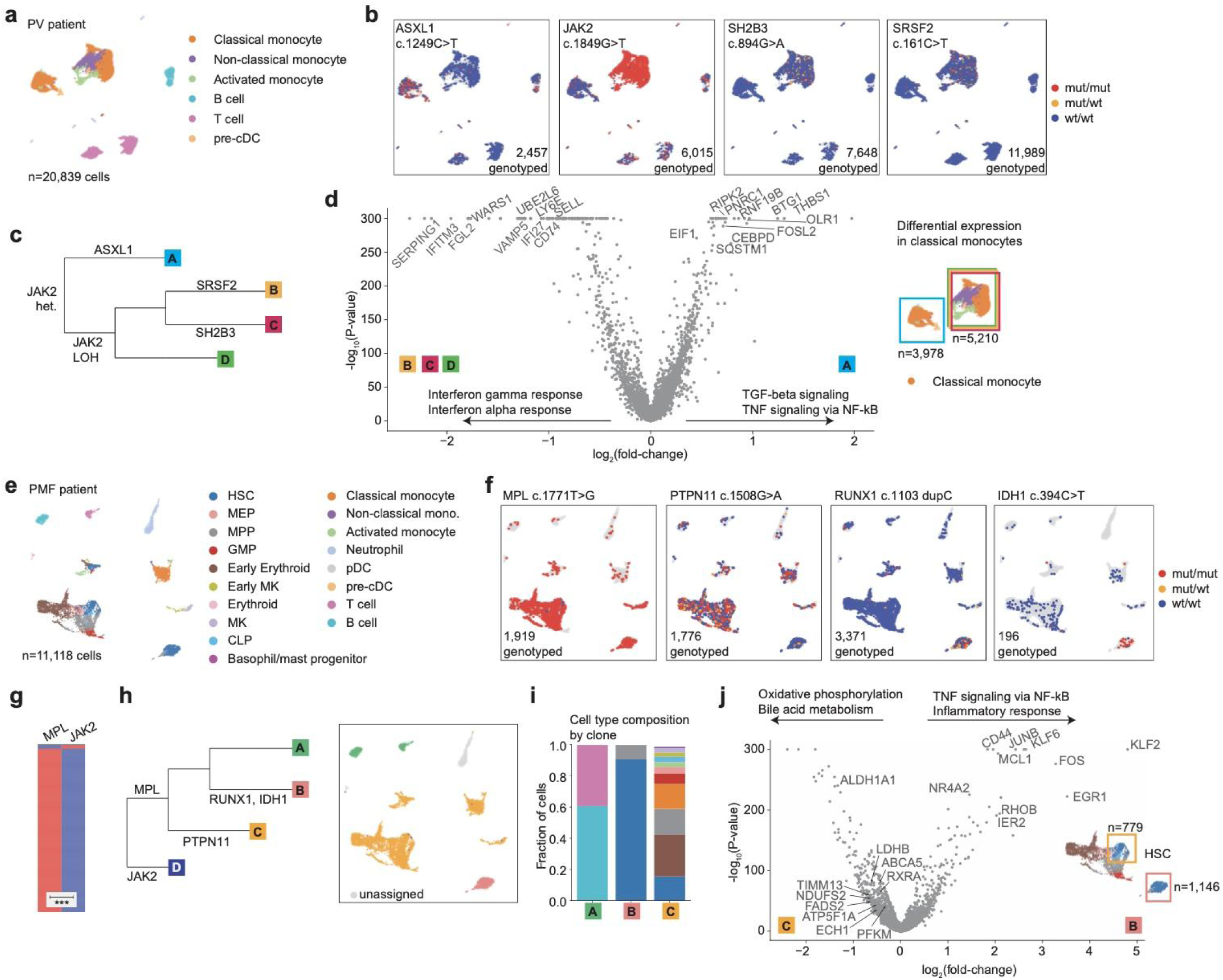
Lineage tracing of additional MPN patients. **a-d**, Genotyping and lineage tracing of an individual patient in the MPN cohort with polycythemia vera (PV). **a-b**, UMAP embedding of this individual patient with cells colored by cell type (**a**) or GIFT genotype for 4 different variants (**b**). **c**, Inferred phylogeny using PICASSO^44^. LOH is inferred by mutation frequencies in each clone. **d**, *Left:* Volcano plot showing differential expression^61^ in classical monocytes of clone A vs. clones B-D (n = 12,066 genes). Significantly enriched pathways are shown for each clone with top pathway genes labeled (FDR < 0.1, methods)^62^. *Right*: Region of UMAP used for differential expression analysis is shown. Only classical monocytes were included in differential expression analysis. **e-j**, Genotyping and lineage tracing of an individual patient in the MPN cohort with primary myelofibrosis (PMF). **e-f**, UMAP embedding of this patient with cells colored by cell type (**e**) or GIFT genotype for 4 different variants (**f**). **g**, Heatmap showing mutually exclusive mutation of *MPL* and *JAK2*, which allows us to separate these mutations into separate clones (Fisher’s exact test, p = 1.3*10^-13^; n = 317 cells genotyped for both variants). **h**, *Left:* Inferred phylogeny using PICASSO^44^ and manual annotation. We included *MPL*, *RUNX1*, and *PTPN11* mutations for PICASSO (n = 275 cells genotyped for all 3 variants), and then we manually added *JAK2* and *IDH1* mutations based on mutation co-occurrence (Fisher’s exact test, as in **g**). *Right:* UMAP embedding colored by clone. Grey indicates wildtype or undetermined clone. **i**, Cell type composition by clone. Colors correspond to the legend in **e**. **j**, *Left:* Volcano plots showing differential expression^61^ in HSCs of clone B versus clone C (n = 12,604 genes). The most significantly enriched pathways are shown for each clone with top pathway genes labeled (FDR < 0.1, methods)^62^. *Bottom Right*: Cells used for differential expression analysis are indicated by boxes on UMAP.

## EXTENDED DATA TABLE LEGENDS

**Extended Data Table 1: Custom probe panels.** Probe names, sequences (lhs_probe, rhs_probe), gapfills (ref_gapfill, alt_gapfill), target types (variant/non-variant/control), probe barcodes (for Flex multiplexing; Visium HD does not use probe barcode), and probe types (gapfill/dual_LHS/dual_RHS/single/control) for all custom probe panels used in this study. **a**, Panel used for cell line mix in Fig. 1. **b**, 4-plex panel spanning a range of gap lengths (Fig. 1f**, Supplementary** Fig. 2i). **c**, Panel used for Visium HD cell line line mix in Fig. 1i**,j**. This panel is the same as **a** but designed for Visium HD capture. **d**, Large multiplexing panel used with four cell lines (Fig. 2a**-d**). **e-f**, Dual probe panel with variant on the LHS (**e**) or RHS (**f**) probe, as shown in Fig. 2e**-g** (Experiment 1 in **Supplementary** Fig. 4c). We included barcodes flanking the LHS and RHS probes to ensure specificity. Less than 1% of UMIs had incorrect barcodes; these were removed from analysis. **g**, 3-plex panel used to directly compare GIFT (gapfill) to dual probes and single probe genotyping (Experiment 2 in **Supplementary** Fig. 4c). **h**, 4-plex panel for profiling glioma (Fig. 3, **Supplementary** Fig. 5). All probe barcodes include the same probes. **i-j**, Probes panels used for profiling MPNs (Fig. 4**-6**). Specific panels used for each patient are listed in **Extended Data Table 2**. **i**, 4-plex panel. All probe barcodes include the same probes. **j**, 16-plex panel. Probes vary slightly by barcode to avoid overlaps.

**Extended Data Table 2: Metadata for glioma and MPN patients in this study. a**, Diagnosis and bulk mutations for glioma patients 1-3 (Fig. 3**, Supplementary** Fig. 5). Mutations were assayed by MSK IMPACT^23^. **b**, Diagnosis and bulk mutations for MPN patients (Fig. 4**-6**). Bulk mutations were assayed from whole blood unless otherwise noted (CD34+ VAFs column). Patient age, sex, and treatments are noted. The GIFT panel used for each patient is also noted (columns “16-plex 1”, “16-plex 2”, “4-plex”, “1-plex”, which correspond to panels in **Extended Data Table 1i-j**). The 1-plex panel included genotyping probes for only *JAK2* V617F. For select patients highlighted in this work, the identification name used in the paper is noted, and/or the figure panel containing phylogenetic analysis for that patient is noted.

**Extended Data Table 3: Gene expression markers for cell typing of MPNs.** Relevant marker genes are listed for MPN cell types.

## Notes

https://github.com/clareaulab/giftwrap

### Summary of Updates

updated license to comply with HHMI funding.

